# Impact of Proximal Tubule-Specific Deletion of Dipeptidyl Peptidase 4 on Blood Pressure, Renal Sodium Handling, and NHE3 Phosphorylation

**DOI:** 10.1101/2024.12.22.629982

**Authors:** Flavia L. Martins, Joao Carlos Ribeiro-Silva, Erika Fernandes de Jesus, Ravi Nistala, Adriana C. C. Girardi

## Abstract

Dipeptidyl peptidase 4 (DPP4) is a transmembrane serine exopeptidase abundantly expressed in the kidneys, predominantly in the proximal tubule (PT); however, its non-enzymatic functions in this nephron segment remain poorly understood. While DPP4 physically associates with the Na^+^/H^+^ exchanger isoform 3 (NHE3) and its inhibitors exert natriuretic effects, the DPP4 role in blood pressure (BP) regulation remains controversial. This study investigated the effects of PT-specific *Dpp4* deletion (*Dpp4*^ΔPT^) and global *Dpp4* deletion (*Dpp4*^−/−^) on systolic blood pressure (SBP), natriuresis, and NHE3 regulation under baseline and angiotensin II (Ang II)-stimulated conditions in both male and female mice. Global and PT-specific *Dpp4* deletion increased diuretic and natriuretic responses to acute saline loading, correlating with enhanced phosphorylation of NHE3 at serine 552 (pS552-NHE3). However, baseline SBP remained unchanged. Ang II stimulation increased DPP4 activity in control mice, with a greater effect in males than in females, reflecting sex-dependent regulation of renal DPP4. In *Dpp4*^ΔPT^ mice, residual kidney DPP4 was unresponsive to Ang II, indicating that PT DPP4, rather than DPP4 in other nephron segments, is regulated by Ang II. Ang II administration increased SBP in all groups; however, the pressor response was significantly attenuated in both *Dpp4*^ΔPT^ and *Dpp4*^−/−^ mice, coinciding with sustained elevated levels of pS552-NHE3. Collectively, these findings demonstrate that PT DPP4 modulates NHE3 activity through mechanisms that prevent the accumulation of pS552-NHE3, exerting an anti-natriuretic effect. In the absence of DPP4, these mechanisms are disrupted, reducing Ang II sensitivity and maintaining high pS552-NHE3 levels, underscoring the role of DPP4 in PT signaling and function.

## INTRODUCTION

Dipeptidyl peptidase 4 (DPP4/CD26) is a widely expressed serine protease found in epithelial and non-epithelial cells across various tissues, with particularly high levels in the kidney^1^. In renal tissue, DPP4 is localized in the glomeruli and the proximal tubule (PT), where it is a major component of the microvilli brush border^2–4^. In addition to its enzymatic activity, DPP4 is involved in a variety of biochemical pathways and physically associates with multiple proteins, including adenosine deaminase^5^, caveolin^6^, components of the extracellular matrix^7,8^, and the sodium-hydrogen exchanger 3 (NHE3)^3,9^.

In the PT, NHE3 mediates approximately 70% of filtered sodium reabsorption, playing a crucial role in extracellular volume homeostasis and blood pressure (BP) control^10–12^. Mice with PT-specific deletion of *Nhe3* display lower BP, enhanced pressure-natriuresis, and attenuated hypertensive responses to chronic angiotensin II (Ang II) infusion compared to wild-type controls^12,13^. Notably, studies have shown that following the onset of hypertension, PT NHE3-mediated sodium reabsorption declines^14–16^, thereby limiting further BP increases^17,18^. This reduction in NHE3 activity is thought to result from increased phosphorylation at serine 552, along with a redistribution of NHE3 from the body to the base of the PT microvilli^14,19^.

Previous work demonstrates that DPP4 inhibitors (DPP4is) downregulate PT NHE3 activity, leading to natriuresis^20–22^. However, despite their natriuretic properties, the impact of DPP4is on BP remains inconclusive. While some studies reported BP reductions in individuals with mild hypertension^23^, chronic kidney disease models^24^, and pre-hypertensive spontaneously hypertensive rats (SHRs)^25^, findings in adult hypertensive animals have been mixed, with outcomes ranging from BP reduction to no change or even BP increases^25–27^.

Given the limited understanding of the physiological role of PT DPP4 and the variable BP responses to DPP4i across different contexts, we generated mice with PT-specific deletion of *Dpp4* and assessed BP, the response to acute saline loading, and phosphorylation of renal NHE3 at serine 552 under both baseline and Ang II-induced BP elevation conditions. To further clarify the specific contribution of PT DPP4, we conducted parallel experiments in global *Dpp4* knockout mice. Additionally, we examined potential sex differences in these regulatory mechanisms.

## METHODS

The data supporting this study’s findings are available from the corresponding authors upon request.

An expanded Methods section is available in the Online-Only Data Supplement.

### Experimental animals

All animal procedures were approved by the Institutional Animal Care and Use Committee of the University of Missouri and the University of São Paulo Medical School in compliance with the National Institutes of Health Guide for the Care and Use of Laboratory Animals. Mice were housed under a 12-hour light/dark cycle in standard rodent cages with free access to standard chow and tap water. Homozygous PT-*Dpp4* knockout mice were generated by crossing *Dpp4*-floxed (*Dpp4*^Fl/Fl^) (Model 10935, Taconic Biosciences, Rensselaer, NY)^28^ female mice with male *Sglt2*-Cre mice^29^ (kindly provided by Dr. Jia L Zhuo, University of Mississippi Medical Center, Jackson, MS, USA). Genotyping was conducted following established protocols (Figure S1 and Table S1). Twelve-week-old mice, including PT-*Dpp4* knockout mice (*Dpp4*^ΔPT^, n=22), *Sglt2*-Cre^negative^*Dpp4*^Fl/Fl^ littermate controls (CTRL, n=22), global *Dpp4* knockout mice (*Dpp4*^−/−^, n=32), and wild-type mice (n=33) were used in this study. Systolic blood pressure (SBP) was measured in acclimated mice using plethysmography, and a saline challenge protocol was conducted by administering an intraperitoneal injection of warmed (37°C) saline (0.9% NaCl) equivalent to 10% of their body weight (v/w)^30^. Immediately after, they were placed in metabolic cages (Tecniplast, Buguggiate, VA, Italy) for a 5-hour urine collection. Urinary volume and sodium excretion were expressed as the percentage of injected fluid and sodium load. To assess the pressor response to Ang II, SBP was recorded 15 minutes before (baseline) and 45 minutes after intraperitoneal Ang II injection (60 µg/kg) (Figure S2). Saline-injected animals served as controls. Kidneys were collected one-hour post-injection, coinciding with peak kidney DPP4 activity and pS552-NHE3 levels (Figure S3). At this time, mice were sedated (4% isoflurane) and subsequently euthanized by cervical dislocation.

### Statistical analysis

Data are presented as mean ± standard error of the mean (SEM). The sample size (n) for each analysis is indicated by individual points in the scatter-dot plots. Comparisons were made using two-way ANOVA followed by Tukey’s post hoc test, with statistical significance set at *P* < 0.05.

## RESULTS

### Phenotypic characterization of PT-specific *Dpp4* deletion in mice

Mice with PT-specific deletion of *Dpp4* (*Dpp4*^ΔPT^) showed a ∼35% reduction in kidney DPP4 in males and a ∼45% reduction in females compared to CTRL mice (Figure 1A). Immunostaining of kidney sections for DPP4 and SGLT2, a PT marker, confirmed that this reduction was specific to the PT. In CTRL mice, DPP4 is evidenced in both the PT, where it colocalizes with SGLT2, and the glomeruli. In contrast, *Dpp4*^ΔPT^ mice showed DPP4 staining exclusively in the glomeruli (Figure 1C). Similarly, kidney DPP4 activity decreased by approximately 30% in *Dpp4*^ΔPT^ males and 40% in *Dpp4*^ΔPT^ females compared to CTRL mice (Figure 1E). Mice with global *Dpp4* deletion (*Dpp4*^−/−^) showed absence of DPP4 protein (Figure 1B), staining (Figure 1D), and activity (Figure 1F). Consistent with previous findings^31^, kidney DPP4 exhibited sexual dimorphism, with higher abundance and activity in females than in males (Figure 1).

**Figure 1.**
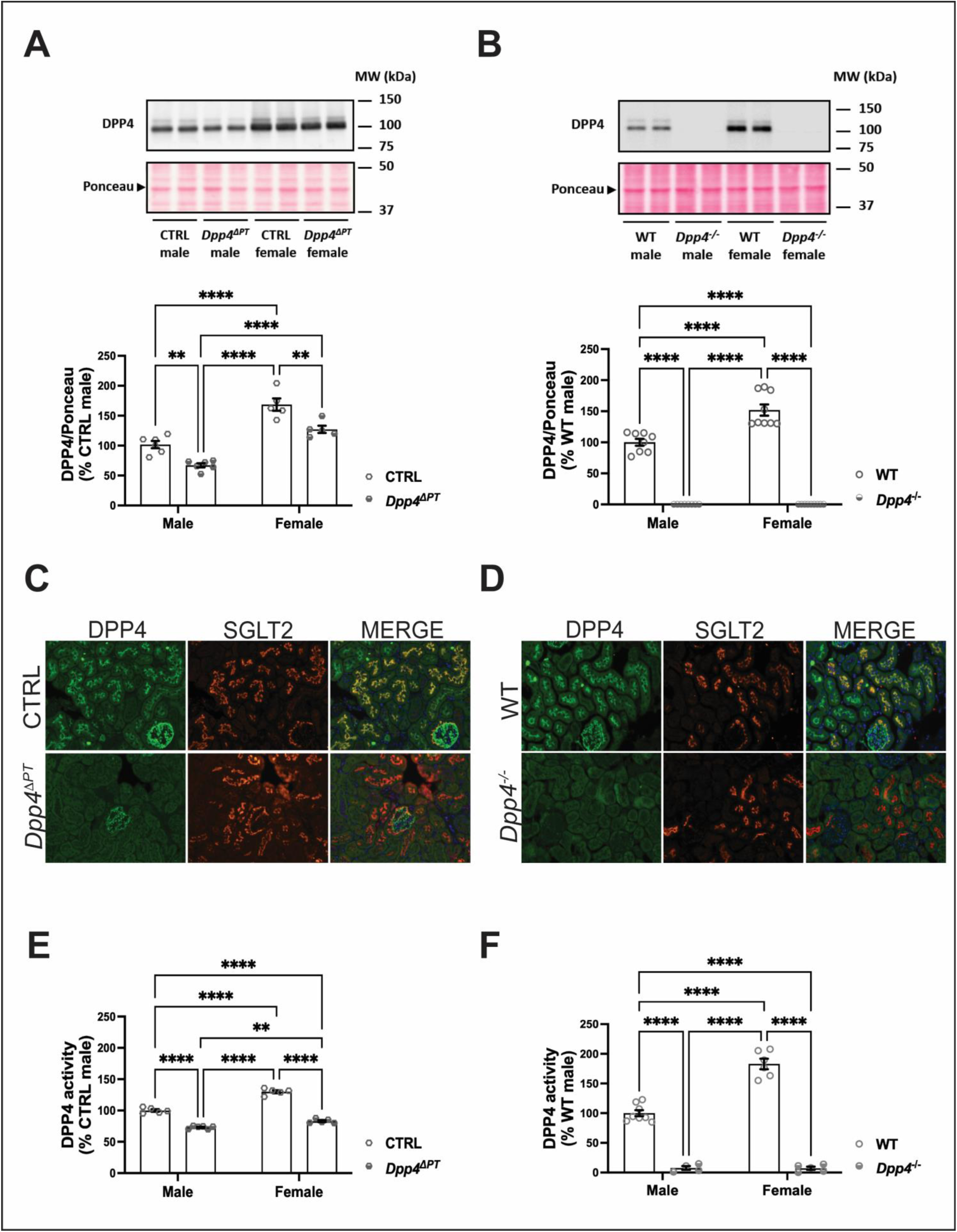
Phenotypic characterization of *Dpp4*^ΔPT^ and *Dpp4*^−/−^ mice. DPP4 protein abundance was evaluated by immunoblotting using equivalent amounts of 10 μg of renal homogenate samples from mice with either **(A)** PT-specific (*Dpp4*^ΔPT^) or **(B)** global (*Dpp4*^−/−^) *Dpp4* deletion and their respective controls. Each dot represents the % of DPP4 expression relative to male CTRL or WT per animal. Representative images of the immunostaining of kidney sections for SGLT2, a PT marker, and DPP4 in **(C)** *Dpp4*^ΔPT^ and **(D)** *Dpp4*^−/−^ mice. Renal DPP4 activity assessed by fluorimetry in renal homogenates from **(E)** *Dpp4*^ΔPT^ and **(F)** *Dpp4*^−/−^ mice. Each dot represents the % of DPP4 activity relative to male CTRL or WT per animal. Bars represent mean ± SEM. **P < 0.01 and ****P < 0.0001.

SBP assessment by plethysmography showed no baseline differences between *Dpp4*^ΔPT^ and CTRL (Figure 2A) or between *Dpp4*^−/−^ and WT mice (Figure 2B), with preserved sex-based BP differences, as female *Dpp4*^ΔPT^ exhibited lower BP than males. Despite comparable SBP, both *Dpp4*^ΔPT^ and *Dpp4*^−/−^ mice exhibited more rapid acute diuretic (Figure 2C-D) and natriuretic (Figure 2E-F) responses to a saline challenge compared to littermate controls. Consistent with previous evidence^31^, acute diuretic (Figure 2C-D) and natriuretic (Figure 2E-F) responses to a saline load were faster in female mice than in males. Interestingly, mice with *Dpp4* deletion (both PT-specific and global) exhibited comparable fluid and salt excretion percentages between males and females (Figure 2C-D).

**Figure 2.**
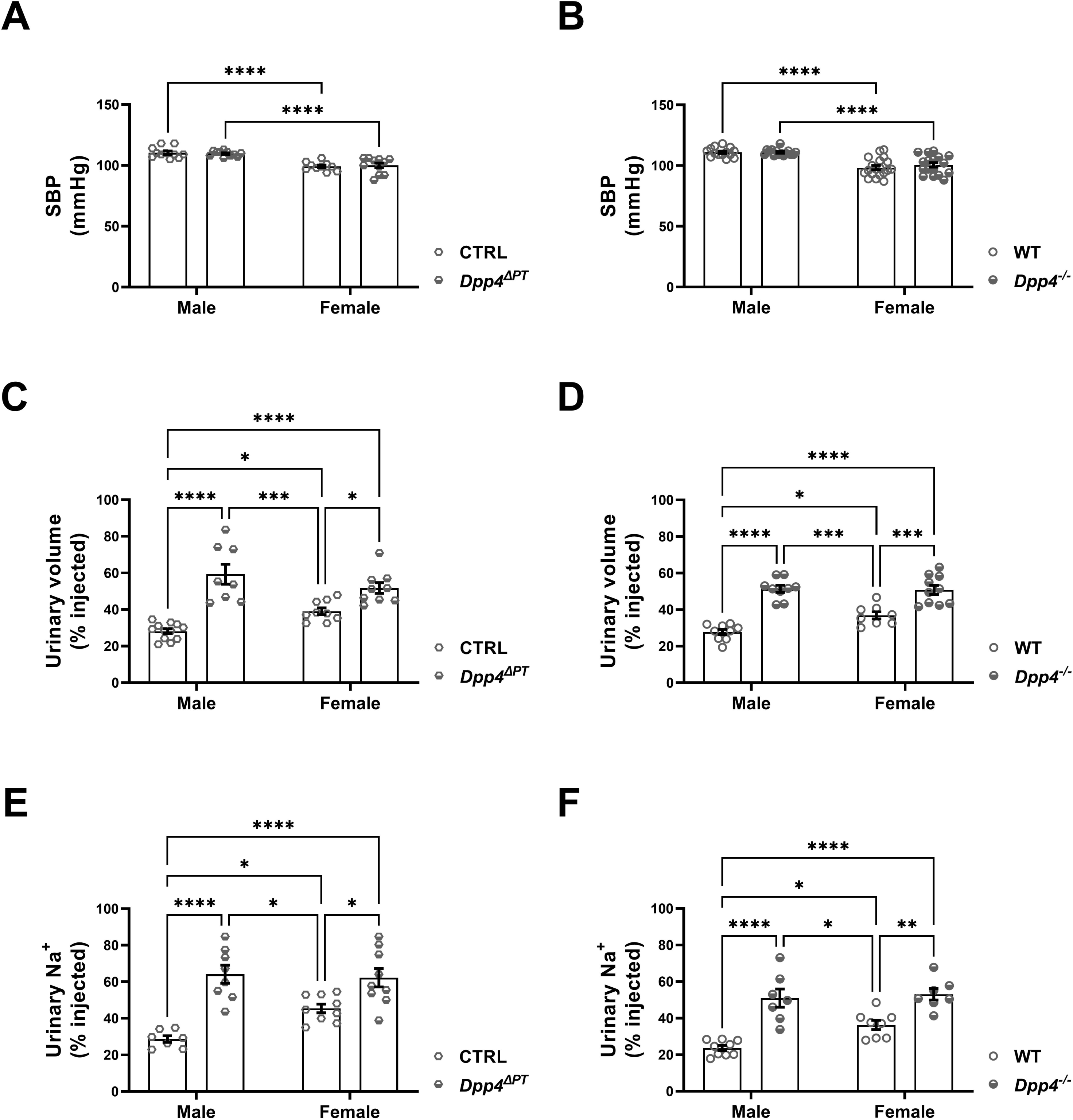
Blood pressure and acute natriuretic and diuretic responses in male and female *Dpp4*^ΔPT^ and *Dpp4*^−/−^ mice. Systolic blood pressure (SBP) was measured by tail-cuff plethysmography in male and female **(A)** *Dpp4*^ΔPT^ and **(B)** *Dpp4*^−/−^ mice. Acute renal natriuretic and diuretic responses were evaluated after a saline challenge. Results expressed as **(C-D)** % of fluid load and **(E-F)** % sodium load excreted within 5 hours. Each dot represents individual measurements. Bars represent mean ± SEM. *P < 0.05, **P < 0.01, ***P < 0.001 and ****P < 0.0001.

The more rapid diuretic and natriuretic responses to a saline challenge in *Dpp4*^ΔPT^ mice suggest reduced sodium and fluid reabsorption in the PT, a function primarily mediated by NHE3. Given that some studies have linked DPP4 inhibition to downregulation of NHE3 activity and increased pS552-NHE3 levels^32^, we investigated kidney pS552-NHE3 levels in our experimental models. CTRL females had higher renal pS552-NHE3 levels than CTRL males, as previously reported^31^. Notably, pS552-NHE3 levels were approximately twofold higher in *Dpp4*^ΔPT^ mice (Figure 3A) and fourfold higher in *Dpp4*^−/−^ mice, in both males and females, compared to their respective controls (Figure 3B). The greater increase in pS552-NHE3 in *Dpp4*^−/−^ mice compared to *Dpp4*^ΔPT^ mice may be partly due to background differences between CTRL (*Dpp4*^Fl/Fl^) and WT mice, as CTRL mice exhibited higher renal pS552-NHE3 levels than WT (Figure S4). Consequently, the difference between *Dpp4*^ΔPT^ and *Dpp4*^Fl/Fl^ was less pronounced than between *Dpp4*^−/−^ and WT.

**Figure 3.**
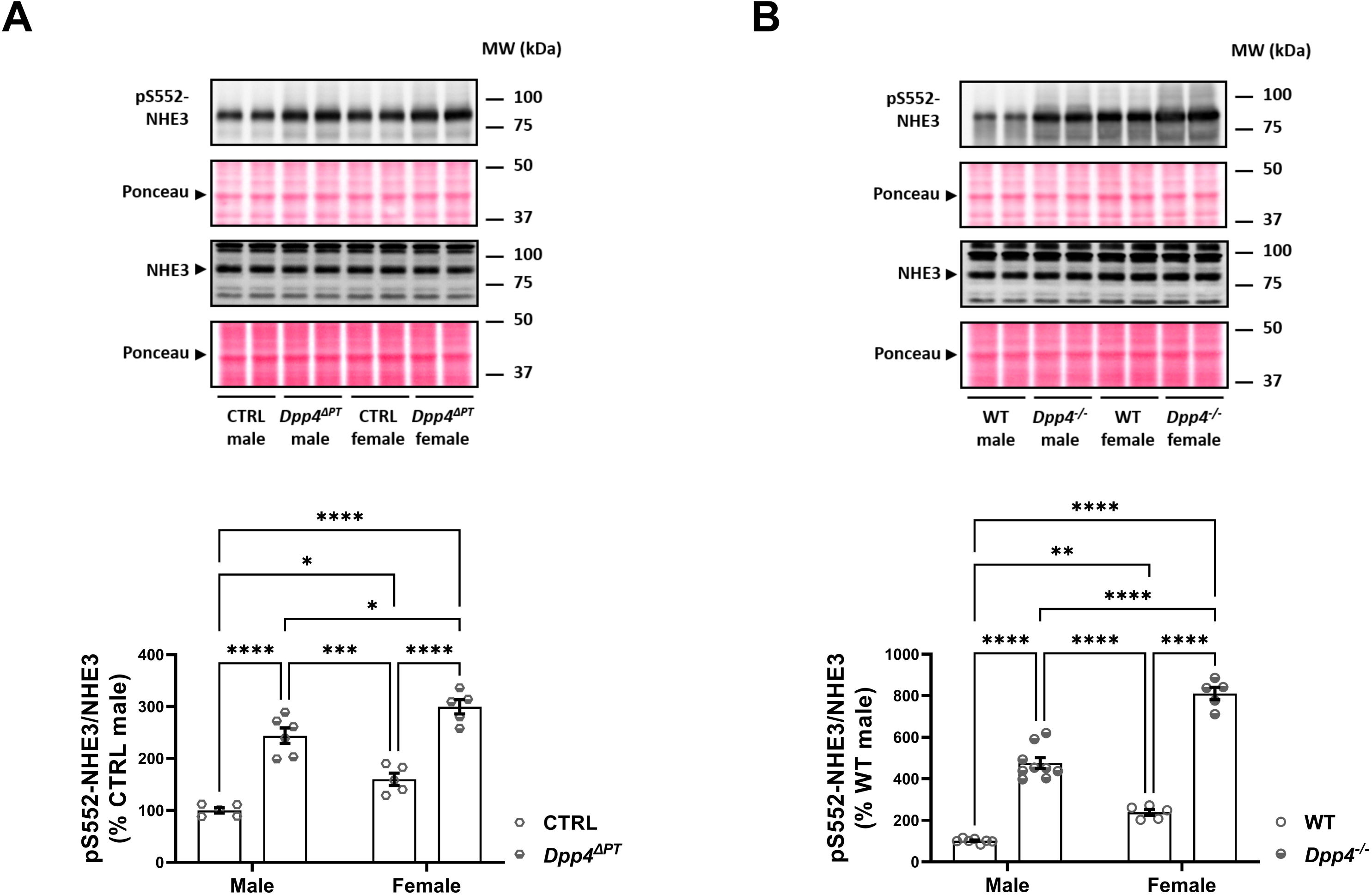
Effect of *Dpp4* deletion on kidney NHE3 (Na^+^/H^+^ exchanger isoform 3) phosphorylation in male and female *Dpp4*^ΔPT^ and *Dpp4*^−/−^ mice. Levels of phosphorylated (pS552-NHE3) and total NHE3 were determined by immunoblotting in kidney homogenates from (A) *Dpp4*^ΔPT^ and (B) *Dpp4*^−/−^ mice. Each dot represents the % of pS552-NHE3/NHE3 relative to male CTRL or WT per animal. Bars represent mean ± SEM. *P < 0.05, **P < 0.01, ***P < 0.001 and ****P < 0.0001.

The sexual dimorphism in pS552-NHE3 was preserved in the absence of DPP4, being predominantly higher in females than male counterparts (Figure 3). The total NHE3 abundance remained unchanged across all experimental groups (Figure S5), consistent with previous findings showing that DPP4 influences NHE3 through posttranslational mechanisms rather than altering its abundance^22,32^.

### Ang II-induced BP elevation is similarly attenuated in both *Dpp4*^Δ^^PT^ and *Dpp4*^−/−^ mice compared to controls

Based on our observations that the absence of DPP4 enhanced the acute diuretic and natriuretic responses, along with elevated pS552-NHE3 in the kidneys of *Dpp4*^ΔPT^ and *Dpp4*^−/−^ mice, we hypothesized that mice lacking *Dpp4* might exhibit an enhanced pressure-natriuresis response, thereby attenuating BP increases. We then investigated whether an acute injection of a pressor dose of Ang II would raise BP to a lesser extent in *Dpp4*^ΔPT^ mice than in CTRL (Figures S2 and S3). Additionally, we examined whether *Dpp4*^−/−^ would have a greater or similar effect on attenuating Ang II-induced BP increases compared to PT-specific deletion. As seen in Figure 4A-B, CTRL mice treated with a supraphysiological concentration of Ang II (60 µg/kg) showed a higher DPP4 activity, with an increase of approximately 50% in males and 30% in females compared to saline. Interestingly, residual kidney DPP4 activity in *Dpp4*^ΔPT^ mice remained unchanged in response to Ang II (Figure 4), suggesting that Ang II specifically regulates PT DPP4 activity. Total DPP4 levels remained unchanged under both saline and Ang II conditions in wild-type mice (Figure S6).

**Figure 4.**
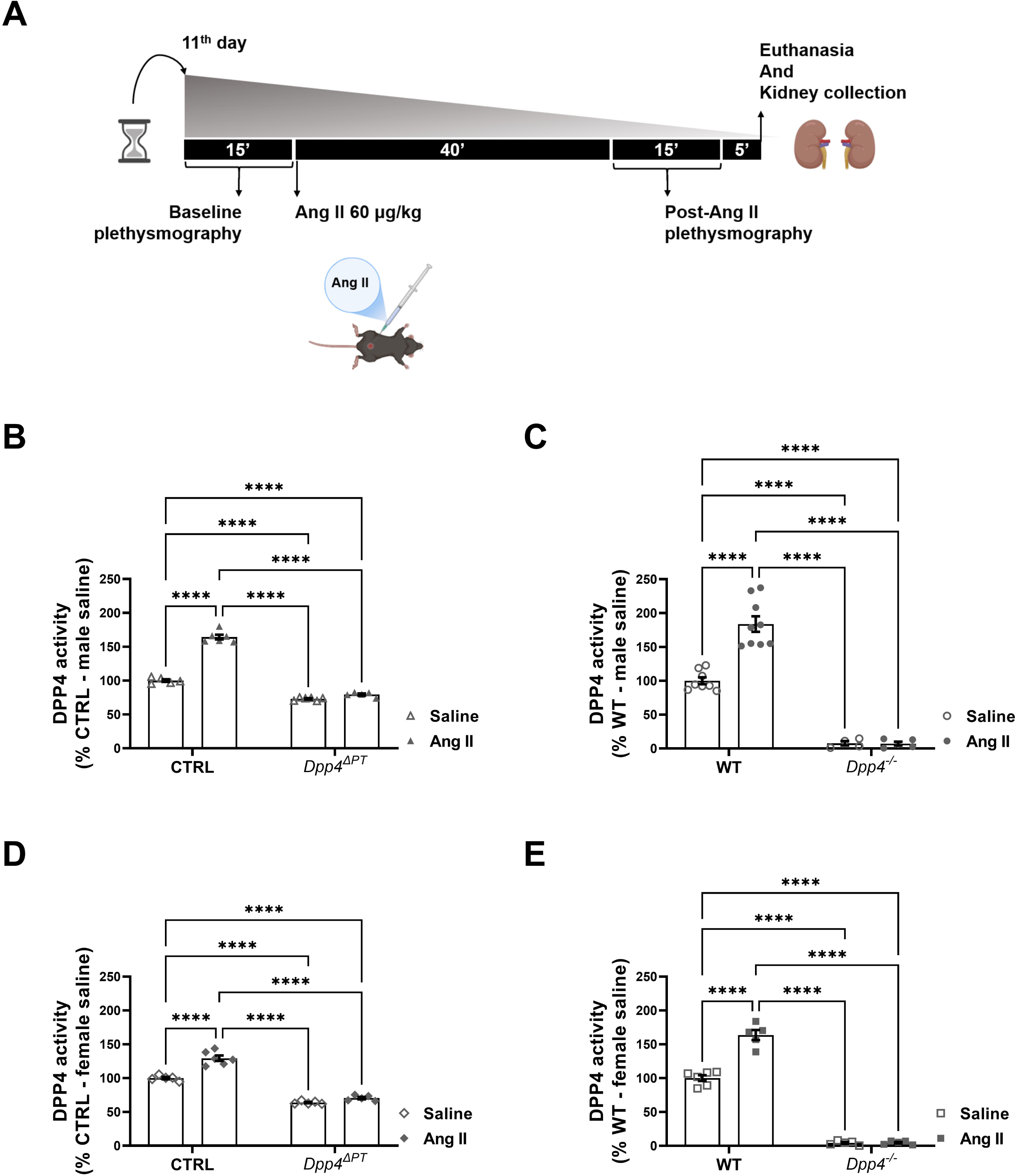
Effect of acute Ang II administration on the renal DPP4 activity of male and female mice. Renal DPP4 activity assessed by fluorimetry in renal homogenates from male and female **(A-B)** *Dpp4*^ΔPT^ and **(C-D)** *Dpp4*^−/−^ mice. Each dot represents the % of DPP4 activity relative to CTRL or WT per animal. Bars represent mean ± SEM. ****P < 0.0001.

SBP was measured before and after Ang II injection (Figure S7), and the change in BP (ΔSBP) was calculated. Ang II administration increased SBP across all experimental groups (Figure S7, right panels). As seen in Figure 5, the pressor response (ΔSBP = Post-Ang II SBP – Baseline BP) was significantly attenuated in *Dpp4*^ΔPT^ compared to CTRL males: 17 ± 1 vs. 29 ± 1 mmHg (P < 0.0001) and females: 20±1 vs. 28 ± 2 mmHg (P < 0.002). Similarly, ΔSBP was also lower in *Dpp4*^−/−^ mice compared to WT males: 24 ± 1 vs. 34 ± 2 mmHg (P < 0.0001) and females: 25 ± 2 vs. 32 ± 3 mmHg (P < 0.03), demonstrating that PT DPP4 contributes to the pressor response to Ang II independently of sex.

**Figure 5.**
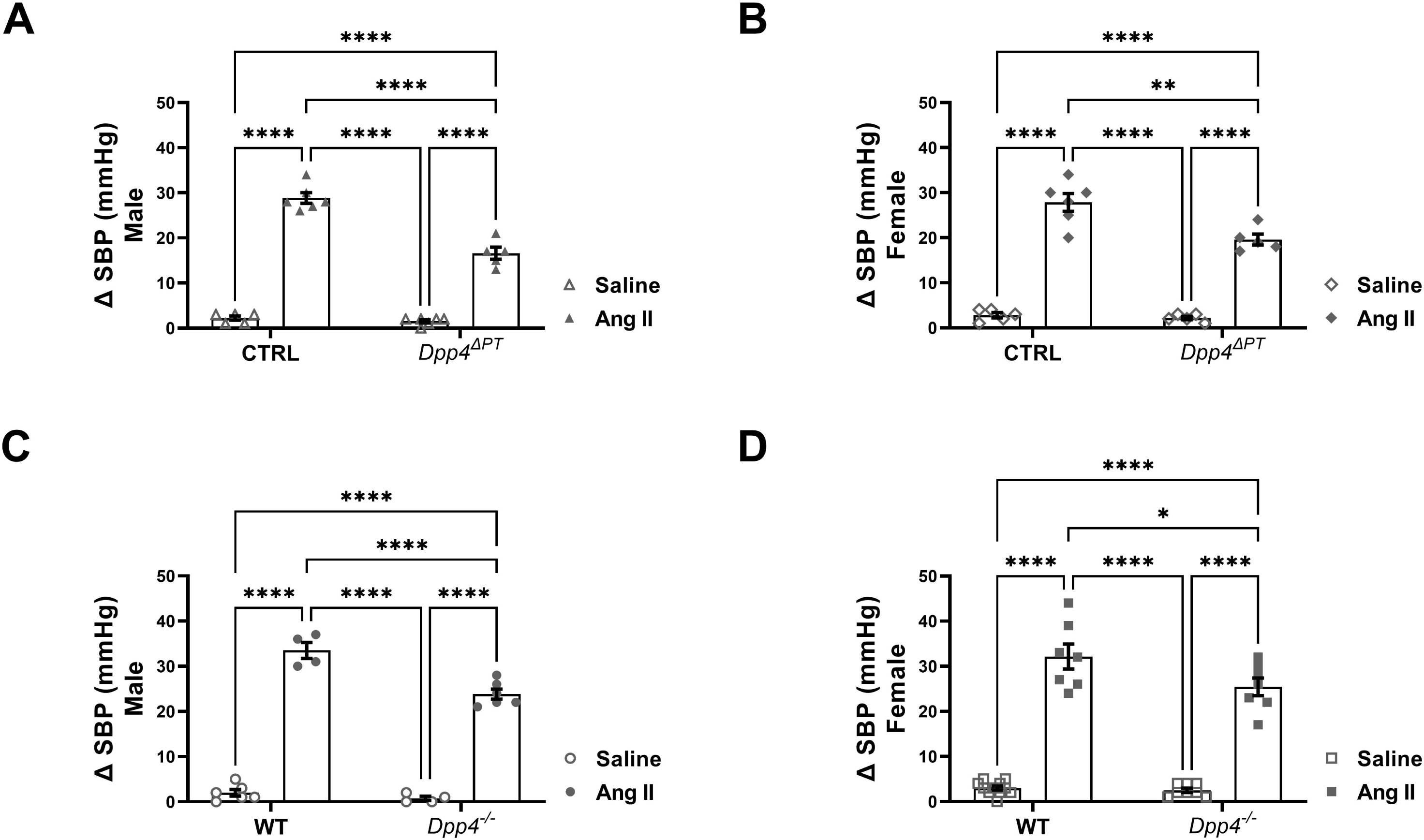
Effect of a pressor dose of Ang II on blood pressure in *Dpp4*^ΔPT^ and *Dpp4*^−/−^ mice. Systolic blood pressure (SBP) was measured by tail-cuff plethysmography before and after Ang II administration in **(A-B)** male and female *Dpp4*^ΔPT^ and **(C-D)** *Dpp4*^−/−^ mice. Each dot represents the ΔSBP change per animal. Bars represent mean ± SEM. *P < 0.05; **P < 0.01 and ****P < 0.0001.

Next, we aimed to determine whether the reduced pressor response to Ang II was associated with further upregulation of pS552-NHE3 in *Dpp4*^ΔPT^ and *Dpp4*^−/−^ mice. In CTRL mice, Ang II administration significantly increased pS552-NHE3 levels (males: Ang II 229 ± 8% vs. saline 100 ± 5%, P < 0.0002; females: Ang II 180 ± 14% vs. saline 100 ± 3%, *P* < 0.0002). In *Dpp4*^ΔPT^ mice, however, Ang II further increased pS552-NHE3 by 95% in males and 61% in females (Figure 6A-B). Similar findings were observed in *Dpp4^−/−^* mice. Ang II increased WT pS552-NHE3 levels (males: Ang II 472 ± 68 vs. saline 100 ± 5%, P < 0.0005; females: Ang II 359 ± 32 vs. saline 100 ± 6%, P < 0.0001). In contrast, Ang II injection in *Dpp4*^−/−^ mice resulted in a greater increase in pS552-NHE3 (176% in males and 104% in females) (Figures 6C and 6D). Total NHE3 levels remained constant across all experimental conditions (Figure S8). Collectively, these findings suggest that the absence of DPP4 enhances NHE3 S552 phosphorylation, thereby attenuating the pressor response to Ang II

**Figure 6.**
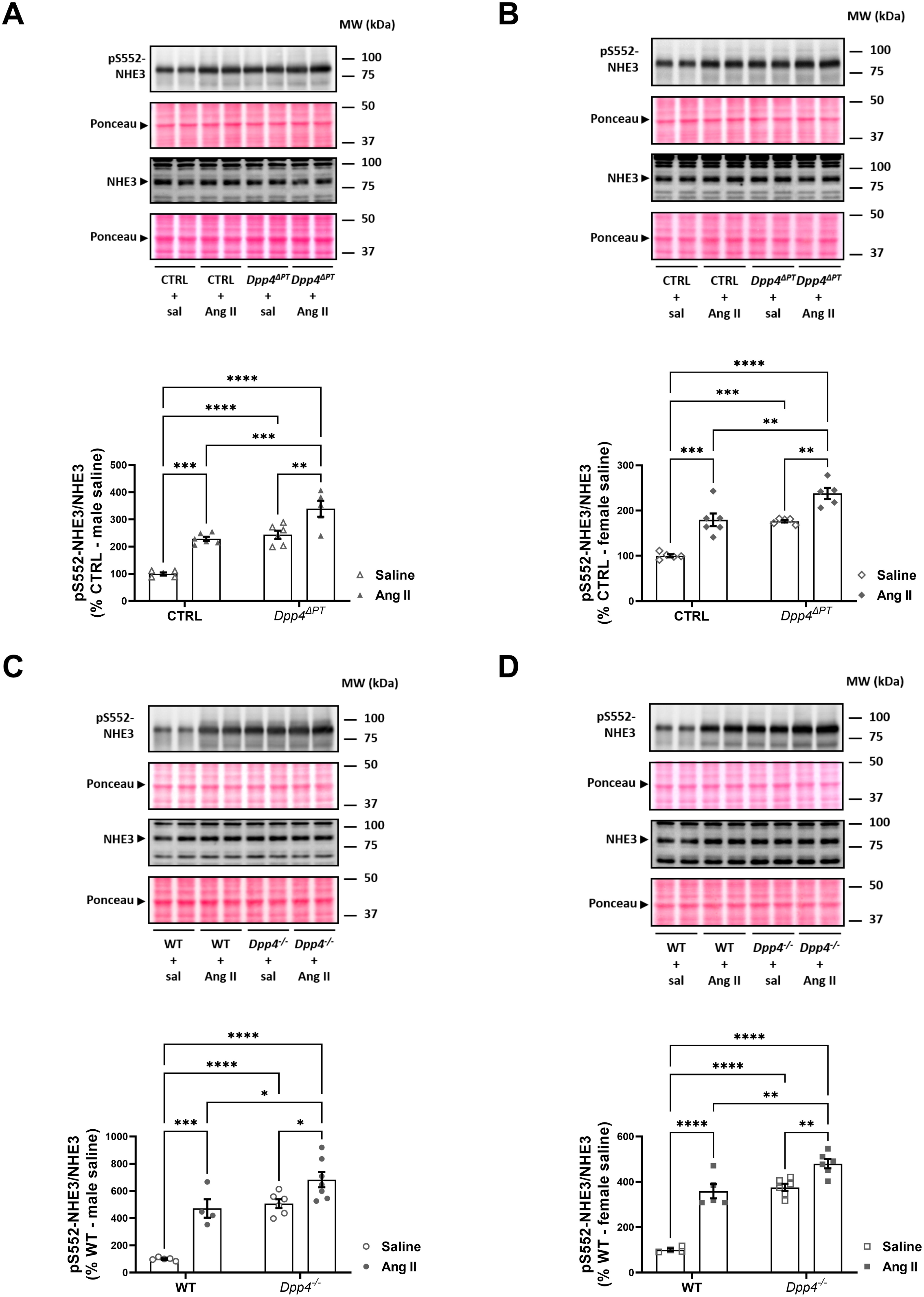
Influence of acute Ang II-induced blood pressure rise on NHE3 phosphorylation in the kidneys of *Dpp4*^ΔPT^ and *Dpp4*^−/−^ mice. Levels of phosphorylated (pS552-NHE3) and total NHE3 were determined by immunoblotting in kidney homogenates from *Dpp4*^ΔPT^ **(A-B)** and *Dpp4*^−/−^ **(C-D)** mice. Each dot represents the % of pS552-NHE3/NHE3 relative to CTRL or WT per animal. Bars represent mean ± SEM. *P < 0.05, **P < 0.01, ***P < 0.001 and ****P < 0.0001.

## DISCUSSION

This study is the first to examine the impact of PT-specific *Dpp4* deletion on natriuresis, renal pS552-NHE3 levels, and BP. Our findings demonstrate that both PT-specific and global *Dpp4* knockout models similarly enhance the natriuretic and diuretic responses to saline load in mice. These findings highlight DPP4’s role in mechanisms regulating salt reabsorption, likely within the PT. Furthermore, both *Dpp4*^ΔPT^ and *Dpp4*^−/−^ mice exhibit upregulation of renal pS552-NHE3 levels, suggesting a baseline reduction in PT NHE3 activity. The comparable reduction in the Ang II-induced BP rise observed in both knockout models, relative to littermate controls, suggests that PT-specific *Dpp4* deletion uniquely counteracts the acute pressor effect of Ang II, likely by enhancing the pressure-natriuresis response.

We previously demonstrated that DPP4 preferentially interacts with NHE3 in the body of the microvilli^3^, where NHE3 is active^33,34^, while phosphorylated NHE3 at serine 552 (pS552-NHE3) localizes to the base of the brush-border microvilli^35^, where NHE3 is inactive^33,34^. In this study, we found that pS552-NHE3 levels are significantly higher in *Dpp4* knockout mice than controls, supporting the notion that baseline NHE3 activity is reduced in the absence of DPP4. These findings raised two key questions: (i) Why does *Dpp4* deletion enhance NHE3 phosphorylation? (ii) Is DPP4 involved in regulating NHE3’s subcellular distribution? As serine 552 (S552) is a consensus site for protein kinase A (PKA)-mediated inhibition of NHE3^36^, one plausible mechanism for increased pS552-NHE3 levels following *Dpp4* deletion is the enhanced bioavailability of DPP4 substrates such as glucagon-like peptide-1 (GLP-1), which activates Gs-coupled receptors^37^. GLP-1 is known to promote natriuresis, at least in part, through PKA-dependent inhibition of NHE3 via pS552 phosphorylation. However, the natriuretic effects of DPP4is are also observed in mice lacking the GLP-1R^21^ and in isolated PT cells^20^ that do not produce GLP-1. These findings suggest that DPP4’s regulation of NHE3 activity and phosphorylation may also occur independently of GLP-1, potentially involving alternative signaling pathways or protein interactions. In this regard, we have previously demonstrated that the interaction between DPP4 and NHE3 is indirect and requires intermediary proteins^38^. Among these, motor proteins involved in NHE3’s subcellular distribution across brush-border microdomains are likely candidates^39^. Ongoing studies aim to clarify these mechanisms and identify additional mediators of the DPP4-NHE3 interaction.

Our data show that female mice exhibit higher DPP4 expression and enzymatic activity, consistent with findings in rats and humans^31,40,41^. Despite DPP4’s role in stimulating NHE3 activity, females paradoxically have higher pS552-NHE3 levels and a faster natriuretic response to saline challenge than males. This discrepancy could be explained by a lower expression of intermediary proteins mediating the DPP4-NHE3 interaction in females, which may reduce NHE3 activation despite elevated DPP4 levels.

Despite elevated levels of renal pS552-NHE3 in both *Dpp4* knockout models, baseline SBP remained unchanged compared to controls, possibly due to compensatory increases in the activity of apical sodium transporters and/or channels in the distal nephron. A potential candidate for this compensation is the sodium-chloride cotransporter (NCC), which we have previously shown to be upregulated in the distal convoluted tubule (DCT) to counteract the inhibition of PT NHE3 by sodium-glucose cotransporter-2 inhibitors (SGLT2i) in normotensive rats^30^. In agreement, we observed that NCC phosphorylated at threonine 53 (pNCC), the active form of NCC^42^, is upregulated in both PT-specific and global *Dpp4* knockout mice (see Supplemental Figure S9). This upregulation likely reflects an adaptive response by the DCT, where increased sodium delivery from PT inhibition stimulates sodium reabsorption capacity in the DCT^43^.

Our findings also demonstrate that the Ang II-mediated BP rise was significantly attenuated in both *Dpp4*^ΔPT^ and *Dpp4*^−/−^. This attenuation was accompanied by further upregulation of kidney NHE3 phosphorylation at serine 552. Elevated kidney pS552-NHE3 and NHE3 redistribution within microvillar microdomains, resulting in reduced NHE3 activity, have been associated with pressure-natriuresis in several hypertension models^14,17,44,45^. In SHRs, for instance, PT NHE3-mediated sodium reabsorption is higher before hypertension onset but subsequently declines compared to normotensive rats^14^. In the pre-hypertensive phase, SHRs show a higher abundance of NHE3 in the body of the microvilli, where it associates with DPP4 and lower pS552-NHE3 levels. Once hypertension is established, however, SHRs this association is reduced and pS552-NHE3 is higher, diminishing PT sodium reabsorption, and contributing to pressure-natriuresis^14^. Similarly, DPP4is attenuate BP in pre-hypertensive SHRs but lose their effectiveness once hypertension is established^25^. A similar pattern is seen in Ang II-induced hypertension, where DPP4is fail to lower BP after hypertension onset^27^. These observations suggest that one plausible explanation for the conflicting data on the effects of DPP4is on BP is that their ability to enhance pS552-NHE3 levels and inhibit NHE3 activity is already maximized in established hypertension, rendering further intervention ineffective. Furthermore, as DPP4is and RAS blockers share overlapping mechanisms^46,47^, their combined use in hypertension therapy warrants further investigation, as it may amplify adverse effects.

Accumulating evidence from our group and others highlights a crosstalk between the signaling pathways activated by Ang II/AT1R and DPP4^40^. In cultured PT cells, supraphysiological concentrations of Ang II enhance DPP4 activity in an ERK 1/2-dependent manner through AT1R activation^48^. Conversely, DPP4is prevent Ang II/AT1R-mediated activation of ERK 1/2. Consistent with these observations, we found that the Ang II-induced increase in DPP4 activity is confined to PT DPP4, as kidney DPP4 activity remained unchanged in *Dpp4*^ΔPT^ mice following Ang II treatment. Interestingly, renal Ang II concentrations were upregulated in Dpp4^ΔPT^ and *Dpp4*^−/−^ (see Supplemental Figure S10), potentially suggesting a compensatory mechanism in response to impaired signaling. Importantly, we have previously shown that the interaction between Ang II/AT1R and DPP4 is pivotal in the pathophysiology of kidney diseases, with DPP4 inhibition preventing glomerular and tubulointerstitial injury, proteinuria, oxidative stress, inflammation, and fibrosis^24,48–50^ processes that are at least partially driven by Ang II/AT1R signaling. Our current findings expand the understanding of the Ang II/AT1R-DPP4 crosstalk, suggesting that it plays a critical role not only in kidney disease pathophysiology but also in proximal tubular function.

In summary, our findings suggest that PT DPP4 exerts an anti-natriuretic effect by tonically stimulating NHE3 through signaling pathways that prevent phosphorylation of serine 552, a key residue associated with the inhibition of PT NHE3-mediated sodium reabsorption. In the absence of DPP4, these regulatory mechanisms are altered, leading to sustained upregulation of pS552-NHE3 levels and reduced BP sensitivity to Ang II, likely due to an enhanced pressure-natriuresis response. Further studies are needed to identify the signaling pathways activated by DPP4 under physiological conditions, as well as their potential impact on NHE3 regulation and other proximal tubular functions.

## SUPPLEMENTAL MATERIAL

Supplemental Methods

Supplementary Tables S1-S2

Figures S1–S17

## NON-STANDARD ABBREVIATIONS AND ACRONYMS

Ang II: Angiotensin II
AT1R: Angiotensin II type 1 receptor
BP: Blood pressure
DPP4: Dipeptidyl peptidase 4
DPP4is: Dipeptidyl peptidase 4 inhibitors
GLP-1: Glucagon-like peptide-1
NHE3: Na^+^/H^+^ exchanger isoform 3
PT: Proximal tubule
RAS: Renin-angiotensin system
SHR: Spontaneously hypertensive rat

