## Supplemental for "Impact of Proximal Tubule-Specific Deletion of Dipeptidyl Peptidase 4 on Blood Pressure, Renal Sodium Handling, and NHE3 Phosphorylation"

### EXPANDED METHODS

*Animals* – All animal procedures were approved by the Institutional Animal Care and Use Committee of the University of Missouri and the University of São Paulo Medical School in compliance with the National Institutes of Health Guide for the Care and Use of Laboratory Animals. All experiments were conducted in 12-week-old male and female mice kept under a 12:12 light/dark cycle, under standard environmental conditions and *ad libitum* access to autoclaved water and standard chow. Heterozygous mice for global *Dpp4* deletion (*Dpp4*$^{+/-}$), were obtained from Taconic Biosciences (Rensselaer, NY). *Dpp4*$^{-/-}$ (homozygous knockout) and *Dpp4*$^{+/+}$ (Wild type, WT) mice were obtained by crossing male and female *Dpp4*$^{+/-}$ mice. *Dpp4* heterozygous floxed mice (*Dpp4*$^{Fl/+}$), kept on a C57BL/6NTac background and backcrossed for at least two generations were obtained from Taconic Biosciences (Model n. 10053) (28) and bred to generate homozygous litters (*Dpp4*$^{Fl/Fl}$). *Dpp4*$^{Fl/Fl}$ mice were then crossed with iL-*Sglt2-Cre* mice (29), (obtained from the Jia L Zhuo lab, University of Mississippi Medical Center) to obtain homozygous PT-*Dpp4* knockout mice.

*Genotyping* – Genotyping conditions were set according to Taconic Biosciences. Briefly, mice tail snips were digested overnight at 55°C or for 1 hour at 85°C under gentle agitation in lysis buffer (Direct PCR - Tail, catalog #102-T, Viagen Biotech, Los Angeles, CA) containing proteinase K (catalog #503-PK, Viagen Biotech). Then, samples were centrifuged at 500 *g* for 5 minutes at 4 °C. For the PCR reaction, buffer composed of 50% DreamTaq Green PCR Master Mix 2x concentrate (catalog #K1081, Thermo Fisher Scientific, Carlsbad, CA), 25 pmol reverse primer, 25 pmol forward primer and 20 ng/ml of DNA sample were mixed and submitted to the program used in the thermal cycler (C1000 Touch Thermal Cycler, BIO-RAD, Hercules, CA). PCR conditions and primer sets (obtained from Integrated DNA Technologies Inc., Coralville, IA) used are described in Supplementary Table S1. Electrophoresis was performed using 1% agarose gel submerged in TAE buffer (40 mM tris acetate pH 8, 10 mM EDTA). SYBR Safe DNA was used for visualization of DNA (catalog # S33102, Invitrogen by Thermo Fisher Scientific). DNA ladder 100 bp (GenDEPOT, Baker, TX) was used as reference. Electrophoresis was performed at 120 V for 25 min. Digitization of the bands was performed using the Odyssey XF Photodocumenter (Li-cor Biotechnology, Lincoln, NB) and ImageStudio software (Li-cor Biotechnology) (Supplementary Figure S1).

*Saline challenge* – Mice were anesthetized with isoflurane (3-4% for induction and 2.5-3% for maintenance) and received an intraperitoneal injection of warmed saline (0.9% NaCl at 37°C) equivalent to 10% of body weight (v/w). Immediately after injection, mice were placed in metabolic cages (Tecniplast, Buguggiate, VA, Italy) and urine over a 5-h period was collected with a graduated pipette. Urinary sodium was measured using chemistry analyzer (AU 480, Beckman Coulter, Brea, CA). Excretion results were presented as the percentage of sodium and fluid load injected.

*Systolic blood pressure (SBP) determination* – SBP was determined with a Hatteras Instruments MC4000 plethysmograph system (Grantsboro, NC). SBP was measured before and 40 minutes after Ang II administration, as previously reported (25) (Supplementary Figure S2). Briefly, mice were acclimated to restraint and tail-cuff inflation for 10 min over six consecutive days. During the experiment, the restrainers' platform was set to heat at 38°C, ensuring sufficient blood flow for acquisition by the equipment, and the first five preliminary measurements were discharged. A total of 10 consecutive and stable readings were considered to calculate the SBP mean at the baseline and after Ang II administration. ΔSBP was determined by subtracting baseline SBP from post-Ang II SBP measurements.

*Acute Ang II administration* – Ang II (catalog #9525, Sigma, St. Louis, MO) was administered intraperitoneally at a dose of 60 µg/kg, and the kidneys were collected 1-hour post-injection for analysis (Supplementary Figure S2), following complete validation in our laboratory (Supplementary Figure S3). Mice were sedated with isoflurane (3-4% for induction and 2.5-3% for maintenance), and the left kidney was immediately excised and placed in ice-cold PBS buffer (150 mM sodium chloride, 2.8 mM monobasic sodium phosphate, 7.2 mM dibasic sodium phosphate, pH 7.4) containing protease inhibitors (0.7 µg/ml pepstatin, 0.5 µg/ml leupeptin and 40 µg/ml phenylmethanesulfonylfluoride) and phosphatase inhibitors (50 mM sodium fluoride and 15 mM sodium pyrophosphate) for subsequent homogenization. The right kidney was fixed in 4% paraformaldehyde in PBS (catalog # J61899, Thermo Fisher Scientific).

*Immunofluorescence Analysis* – The right kidney was fixed in 4% paraformaldehyde in PBS (catalog #J61899, Thermo Fisher Scientific) for 24 hours, stored in 70% ethanol, and then embedded in paraffin. Tissue samples were cut into 4-μm sections and placed on silanized slides (StarFrost, Knittel Glass, Bielefeld, Germany). Sections were deparaffinized, rehydrated, and subjected to antigen retrieval by heating at 140°C for 20 minutes in 10 mM citrate buffer (pH 6.0) supplemented with 1% SDS-PBS. Next, sections were washed twice in deionized water, gently dried, and demarcated using a hydrophobic barrier pen (PAP pen, catalog #AB2601, Abcam, Cambridge, UK). To reduce nonspecific binding, blocking was performed with 5% BSA and 0.25% Triton X-100 in PBS for 30 minutes. Sections were then incubated overnight at 4°C with primary antibodies against DPP4 (1:100, catalog #AF954, R&D Systems, Minneapolis, MN) and SGLT2 (1:100, catalog #20802, Bicell, St. Louis, MO) diluted in the blocking solution. Following primary antibody incubation, sections were washed three times with PBS (5 minutes each). Sections were then incubated with Alexa Fluor 488-conjugated donkey anti-goat IgG (catalog #A11055, Life Technologies/Thermo Fisher Scientific), Alexa Fluor 647-conjugated donkey anti-rabbit IgG (catalog #A31573, Life Technologies/Thermo Fisher Scientific), and DAPI (catalog #62248, Thermo Fisher Scientific) diluted 1:500 in the blocking solution for 1 hour at room temperature. Afterward, sections were washed again in PBS (3 × 5 minutes), mounted in Fluoromount-G (catalog #00-4958-02, Invitrogen/Thermo Fisher Scientific), and coverslipped (Knittel Glass). Finally, sections were allowed to dry at room temperature before visualization with an EVOS M7000 microscope (Invitrogen/Thermo Fisher Scientific). Fluorescent excitation and detection parameters were kept identical across all experimental groups.

*Kidney homogenate preparation* – The left kidney was homogenized in ice-cold PBS containing protease and phosphatase inhibitors using a Potter–Elvehjem-style tissue grinder (POLIMIX PX-SR50E, Kinematica Inc., Luzern, Switzerland). The homogenate was then cleared by centrifugation (2,400 × g for 10 minutes at 4°C), aliquoted, and stored at −80°C. Protein concentration was determined by the bicinchoninic acid (BCA) method using the Pierce BCA Protein Assay Kit (Thermo Fisher Scientific).

*SDS-PAGE and Immunoblotting* – Mouse kidney homogenates were solubilized in Laemmli sample buffer (catalog #1610737, Bio-Rad) and separated using either commercial (Criterion TGX, catalog #5671125, Bio-Rad) or homemade 10% SDS-PAGE gels. Proteins were then transferred overnight at 4°C onto polyvinylidene fluoride membranes (catalog #88518, Thermo Fisher Scientific) using the Criterion Blotter with Wire Electrodes (catalog #17704071, Bio-Rad) at 30 V for 16 hours at 4°C. Membranes were blocked for 1 hour in PBS containing 5% nonfat dry milk and 0.1% Tween-20, then incubated overnight at 4°C with primary antibodies (see Supplementary Table S2 for details). The following day, membranes were washed, incubated for 1 hour at room temperature with the corresponding horseradish peroxidase–conjugated secondary antibodies, and washed again. Proteins were visualized using an enhanced chemiluminescence detection system (catalog #34095, SuperSignal West Femto Maximum Sensitivity Substrate, Thermo Fisher Scientific) on an Odyssey XF photodocumenter (Li-Cor Biotechnology, Lincoln, NE) with ImageStudio software (Li-Cor Biotechnology). Band intensities were quantified using Scion Image software (Scion, Frederick, MD), and results were expressed as a percentage of the control for each gel.

*DPP4 activity assay* – Kidney DPP4 activity was evaluated in 100 μg of renal homogenates diluted in DPP4 assay buffer (Tris-HCl pH 8.0 and 150 mM NaCl). 50 μl of the diluted sample was added to a 96-well black flat-bottom plate. Next, 50 μl of the fluorescent DPP4 substrate, 100 μM H-Ala-Pro-AFC (I-680, Bachem, Torrence, CA), was added to each well and incubated for 10 min at room temperature, protected from light. Fluorescence was measured using the Synergy Microplate Reader (Biotek, Winooski, VT) with an excitation wavelength of 405 nm and an emission wavelength of 535 nm. The assays were conducted in the presence and absence of the DPP4 inhibitor (10 μM linagliptin, Sigma) in duplicates. The result was calculated as relative light units (RLU) and presented as a percentage of the respective control group.

*Renal Ang II content* - Renal angiotensin II content was measured by Enzyme Linked Immuno Sorbent Assay (Biomatik®, Cambridge, ON - catalog #EKU02406) following the manufacturer's instructions.

*Statistical analysis* – Data are presented as mean ± standard error of the mean (SEM). The sample size (n) for each analysis is indicated by individual points in the scatter-dot plots. Statistical analyses were performed using GraphPad Prism 10.0 (San Diego, CA). Normality was assessed with the Shapiro-Wilk test. Group comparisons were conducted using Student's t-test for two groups (Figures S4, S9–10), and one-way (Figure S3) or two-way ANOVA (Figures 1–6, S5–S8) with Tukey's post hoc test for four or more groups. Statistical significance was set at $P < 0.05$.

**Supplementary Table S1.** Genotyping conditions.

| Reaction | Primer sequence | PCR conditions |
|---|---|---|
| *Dpp4*$^{-/-}$ | Forward 5' – GAATATGATCCTTGTCAGAGCAGCC - 3'<br>Reverse 5' – CTGCACTCAGAAGTCTCACTG – 3'<br><br>Control primer<br>Forward 5' – GAGACTCTGGCTACTCATCC – 3'<br>Reverse 5' – CCTTCAGCAAGAGCTGGGGAC – 3' | Step 1: 95°C, 15 min<br>Step 2: 94°C, 45 secs<br>Step 3: 60°C, 1 min<br>Step 4: 72°C, 1min<br>(37 cycles – Steps 2-4)<br>Step 5: 72°C 5min<br>Cooling 4°C ∞ |
| *Dpp4*$^{Fl/Fl}$ | Forward 5' – GGAGGGTGAATTTATAATCCTTTACC - 3'<br>Reverse 5' – CTGCACTCAGAAGTCTCACTG – 3' | Step 1: 95°C, 15 min<br>Step 2: 94°C, 45 secs<br>Step 3: 60°C, 1 min<br>Step 4: 72°C, 1min<br>(35 cycles – Steps 2-4)<br>Step 5: 72°C 5min<br>Cooling 4°C ∞ |
| *Cre* | Forward 5' – CCTGGAAAATGCTTCTGTCCG- 3'<br>Reverse 5' – CAGGGTGTTATAAGCAATCCC – 3' | Step 1: 95°C, 15 min<br>Step 2: 94°C, 45 secs<br>Step 3: 60°C, 1 min<br>Step 4: 72°C, 1min<br>(35 cycles – Steps 2-4)<br>Step 5: 72°C 5min<br>Cooling 4°C ∞ |

**Supplementary Table S2.** Antibody details. IB: immunoblotting, IF: immunofluorescence.

| Target | Method | Species raised | Molecular weight (kDa) | µg protein/lane | 1ARY antibody specifications | 2ARY antibody specifications |
|---|---|---|---|---|---|---|
| **DPP4** | IB | Goat | 110 | 10 | AF954<br>R&D Systems (Minneapolis, MN)<br>1:800 v/v | 305-035-003<br>Jackson Immuno Research<br>Peroxidase AffiniPure™ (HRP) Rabbit Anti-Goat IgG<br>1:2000 v/v |
| **NHE3** | IB | Rabbit | 80 | 20 | Alicia McDonough (USC, Los Angeles, CA)<br>1:2000 v/v | 111-035-144<br>Jackson Immuno Research<br>Peroxidase AffiniPure™ (HRP) Goat Anti-Rabbit IgG<br>1:2000 v/v |
| **pS552-NHE3** | IB | Mouse | 80 | 10 | Sc-53962<br>Santa Cruz Biotechnology (Santa Cruz, CA)<br>1:1000 v/v | 115-035-062<br>Jackson Immuno Research<br>Peroxidase AffiniPure™ (HRP) Goat Anti-Mouse IgG<br>1:2000 v/v |
| **pT53-NCC** | IB | Rabbit | 150 | 10 | P1311-53<br>PhosphoSolutions<br>(Aurora, CO)<br>1:5000 v/v | 111-035-144<br>Jackson Immuno Research<br>Peroxidase AffiniPure™ (HRP) Goat Anti-Rabbit IgG<br>1:2000 v/v |
| **DPP4** | IF | Goat | - | - | AF954<br>R&D Systems (Minneapolis, MN)<br>1:100 v/v | A11055<br>Life Technologies<br>Alexa Fluor™ 488 Donkey anti-goat IgG<br>1:500 v/v |
| **SGLT2** | IF | Rabbit | - | - | 20802<br>Bicell<br>(St. Louis, MO)<br>1:100 v/v | A31573<br>Life Technologies<br>Alexa Fluor™ 647 Donkey anti-rabbit IgG<br>1:500 v/v |

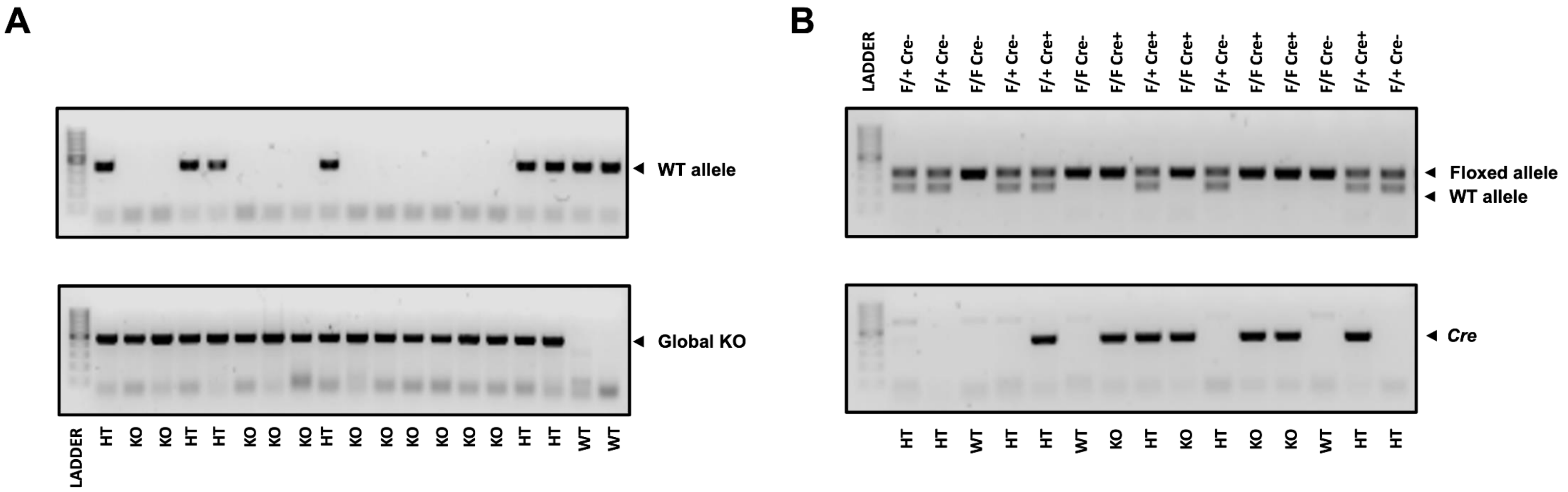


**Supplementary Figure S1 – Genotyping of PT-Specific and global *Dpp4* knockout mice. (A)** Genotyping of global *Dpp4* knockout (*Dpp4*$^{-/-}$) mice. The top reaction was performed to detect the wild-type allele, whereas the bottom reaction was performed to detect the mutated *Dpp4* allele. **(B)** Genotyping of PT-specific *Dpp4* knockout (*Dpp4*$^{\Delta PT}$) mice. The top reaction was performed for detection of the wild type and floxed *Dpp4* alleles, while the bottom reaction was performed to detect the presence of the *Cre* allele.

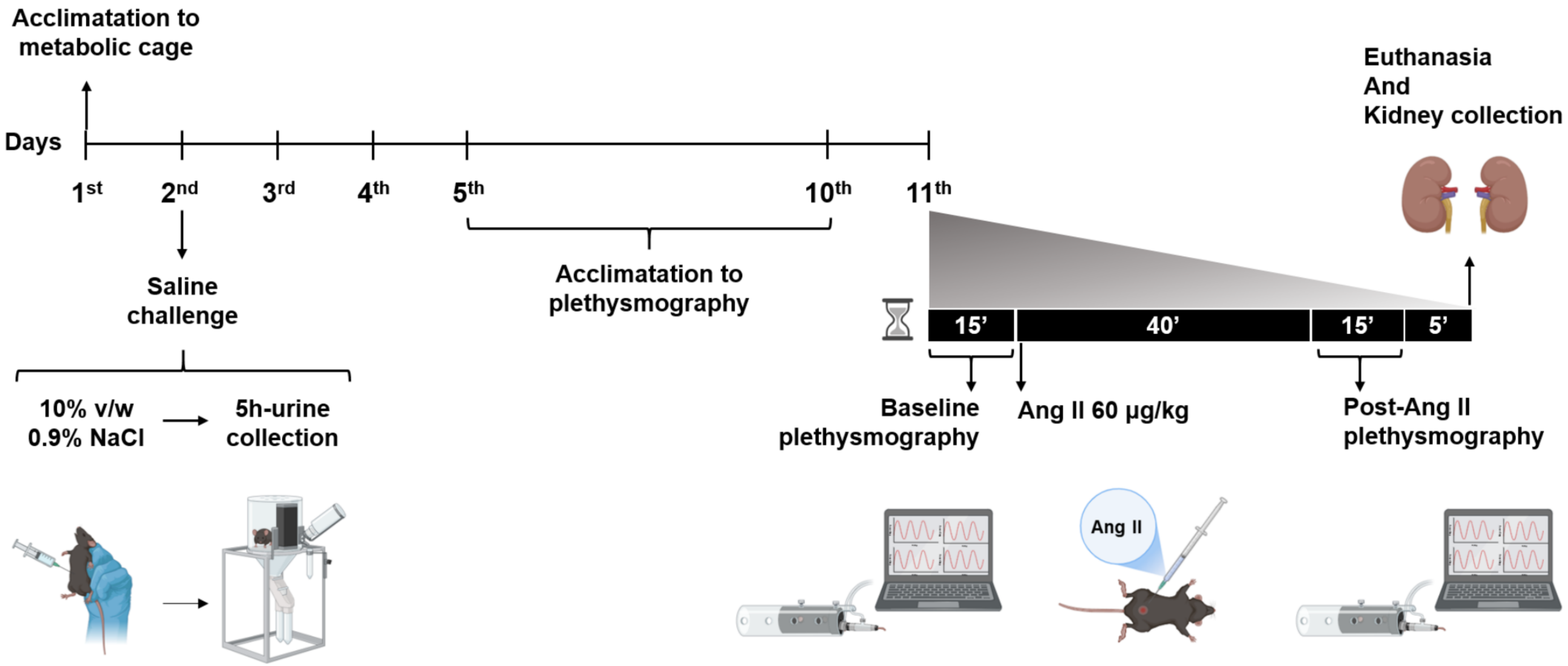


**Supplementary Figure S2. Experimental design –** Experiments were conducted in 12-week-old male and female mice. Following a 24-hour acclimation period in metabolic cages, mice received an intraperitoneal injection of 0.9% NaCl. The next two days served as a rest period with no mice manipulation. Afterward, mice were acclimated for noninvasive systolic blood pressure (BP) measurement via tail-cuff plethysmography over six consecutive mornings, with each session lasting approximately 10 minutes. On the sixth day, SBP was measured and designated as the baseline. Subsequently, mice received an intraperitoneal infusion of Ang II (60 ng/kg) or saline, were returned to their cages for 40 minutes, and then underwent a 15-minute SBP measurement session via tail-cuff plethysmography. Kidneys were collected in ice-cold PBS containing protease and phosphatase inhibitors or fixed in 4% paraformaldehyde for further analysis.

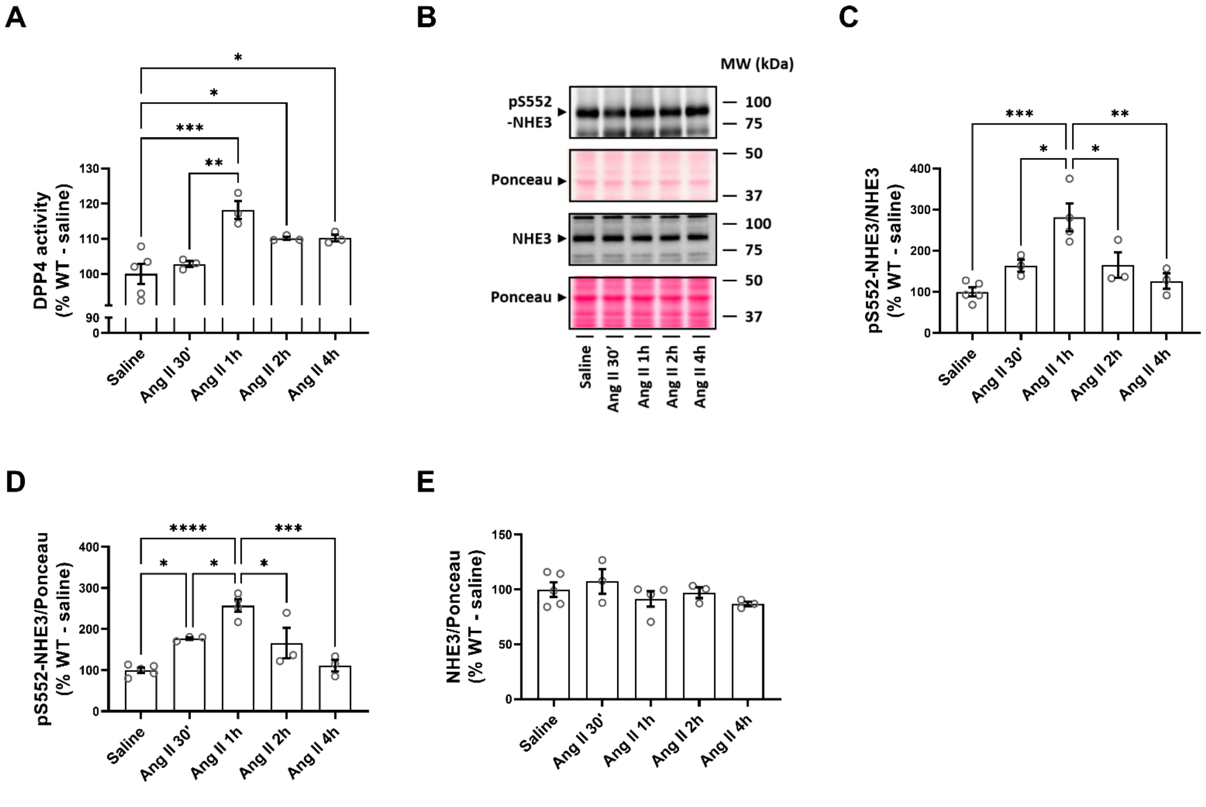


**Supplementary Figure S3 – Standardization of the experimental protocol for acute Ang II administration.** A pressor dose of Ang II (1000 ng/kg/min, equivalent to 60 µg/kg) was administered intraperitoneally to male WT mice. **(A)** DPP4 activity measured by fluorimetry. **(B)** Immunoblot evaluation of pS552-NHE3 and total NHE3 expression. **(C)** pS552-NHE3/NHE3 ratio. **(D)** Quantification of pS552-NHE3 normalized by Ponceau staining. **(E)** Quantification of total NHE3 normalized by Ponceau staining. Values are expressed as a percentage of WT and presented as mean ± SEM. One-way ANOVA followed by Tukey's post-test was used for comparisons. *P < 0.05, **P < 0.01, ***P < 0.001 and ****P < 0.0001.

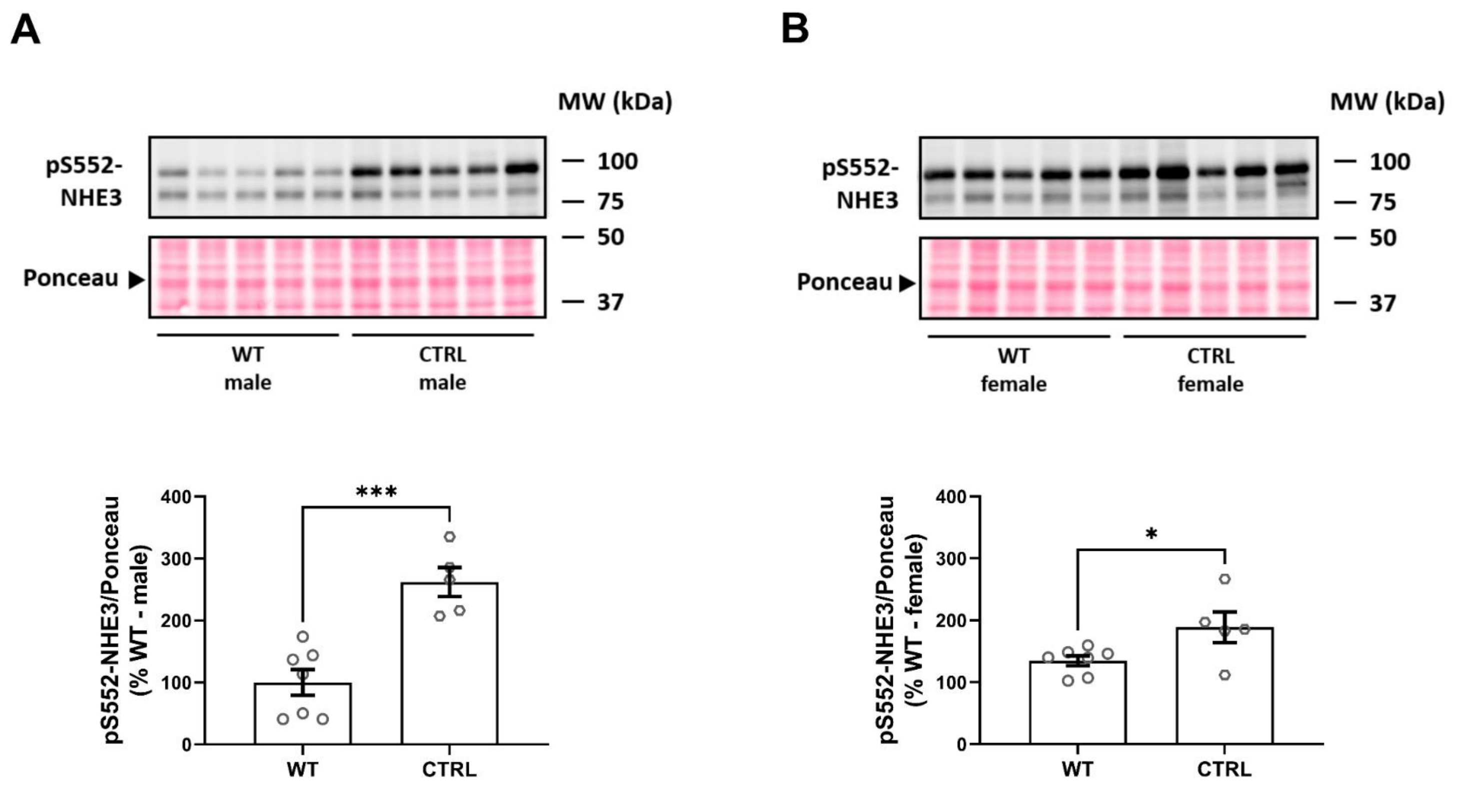


**Supplementary Figure S4 – Baseline differences in NHE3 phosphorylation at serine 552 in male and female CTRL and WT mice. Evaluation of the levels of** NHE3 phosphorylation at serine 552 in kidney homogenates from **(A)** male or **(B)** female CTRL (*Dpp4*$^{\Delta PT}$ littermate) and WT (*Dpp4*$^{-/-}$ littermate) mice. Densitometric analyses were normalized to the Ponceau signal (~42 kDa). Data are expressed as a percentage of the respective WT (male or female) controls and shown as mean ± SEM. Statistical significance was determined by Student's *t*-test. *P < 0.05 and ***P < 0.001.

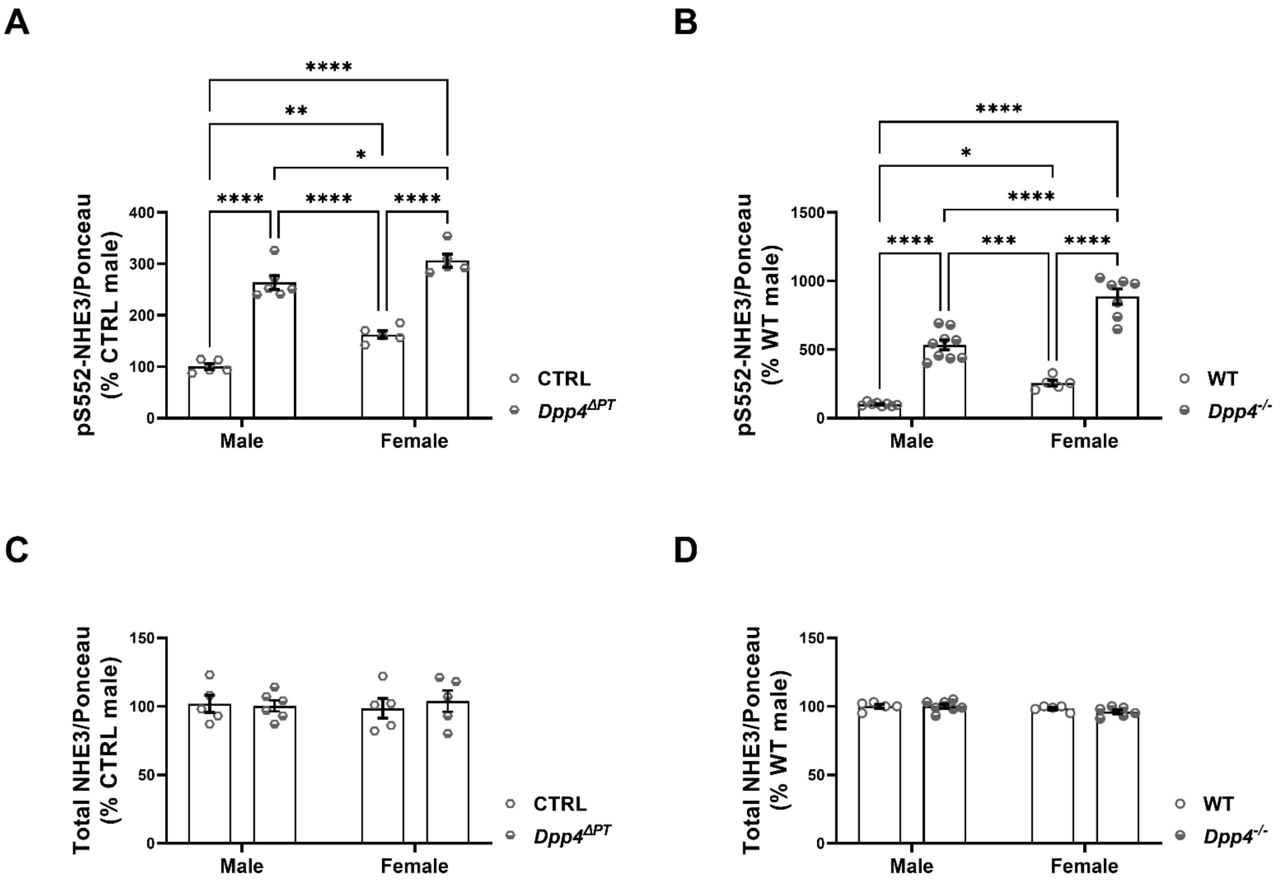


**Supplementary Figure S5 – Effect of PT-specific and global deletion of *Dpp4* on NHE3 phosphorylation at serine 552 and total NHE3 abundance in male and female mice.** Graphical representation of the densitometric analysis of the immunoblotting performed with antibodies against pS552-NHE3 (1:1000) and total NHE3 (1:2000) in kidney homogenates from **(A, C)** *Dpp4*$^{\Delta PT}$ and **(B, D)** *Dpp4*$^{-/-}$ mice. Densitometry results were normalized to Ponceau staining (42 kDa) and expressed as % of CTRL or WT male values. Data expressed as mean ± SEM. Statistical analysis was performed using two-way ANOVA followed by Tukey's post-test. *P < 0.05, **P < 0.01, ***P < 0.001 and ****P < 0.0001.

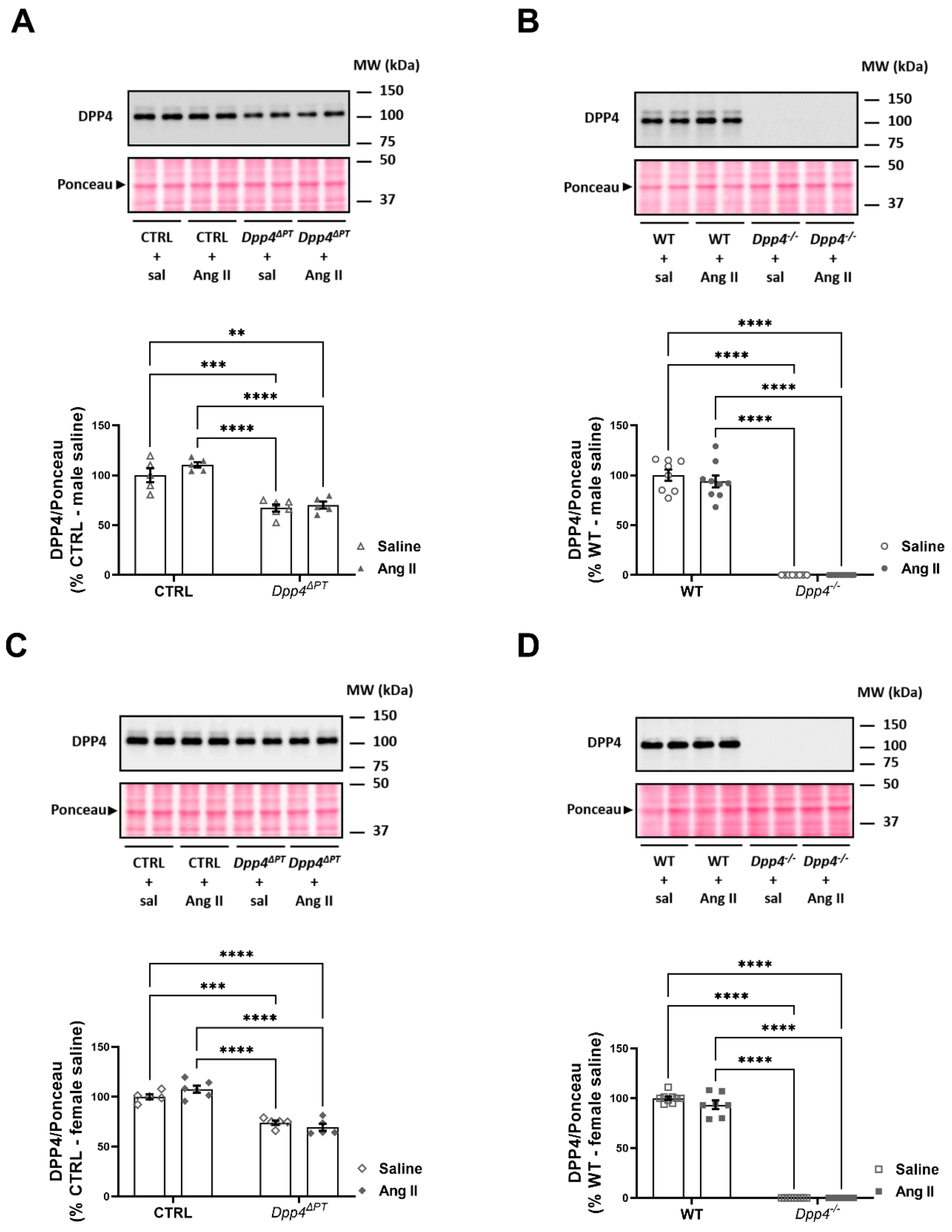


**Supplementary Figure S6 – Effect of acute Ang II administration on renal DPP4 protein abundance in male and female *Dpp4*$^{\Delta PT}$ and *Dpp4*$^{-/-}$ mice.** DPP4 abundance was analyzed by immunoblotting in kidney homogenates (10 µg) from **(A)** male *Dpp4*$^{\Delta PT}$, **(B)** male Dpp4$^{-/-}$, **(C)** female *Dpp4*$^{\Delta PT}$ and **(D)** female *Dpp4*$^{-/-}$ mice. The densitometry results were normalized to Ponceau staining (~42 kDa) and expressed as % of CTRL or WT male/female. Data expressed as mean ± SEM. Statistical analysis was performed using two-way ANOVA followed by Tukey's post-test. ***P < 0.001 and ****P < 0.0001.

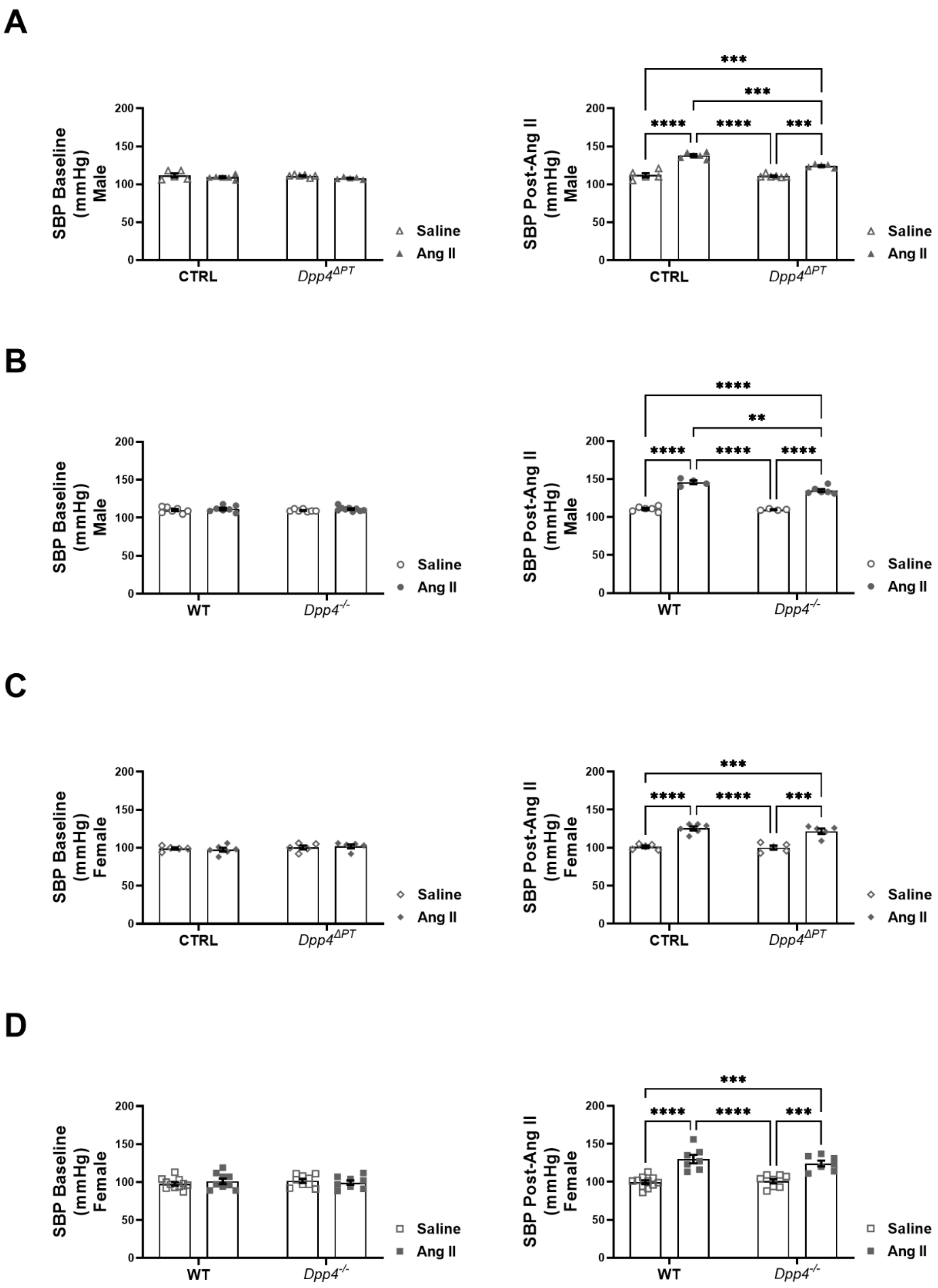


**Supplementary Figure S7 – Baseline and post-Ang II systolic blood pressure (SBP) in male and female *$Dpp4^{\Delta PT}$* and *$Dpp4^{-/-}$* mice.** SBP was measured by tail-cuff plethysmography in *Dpp4*$^{\Delta PT}$, *Dpp4*$^{-/-}$, and their respective male and female littermate controls. Panels **A-D** show baseline SBP (left) and SBP after 1 h of Ang II stimulation (right). Statistical analysis was performed using two-way ANOVA followed by Tukey's post-test. **P < 0.01, ***P < 0.001 and ****P < 0.0001.

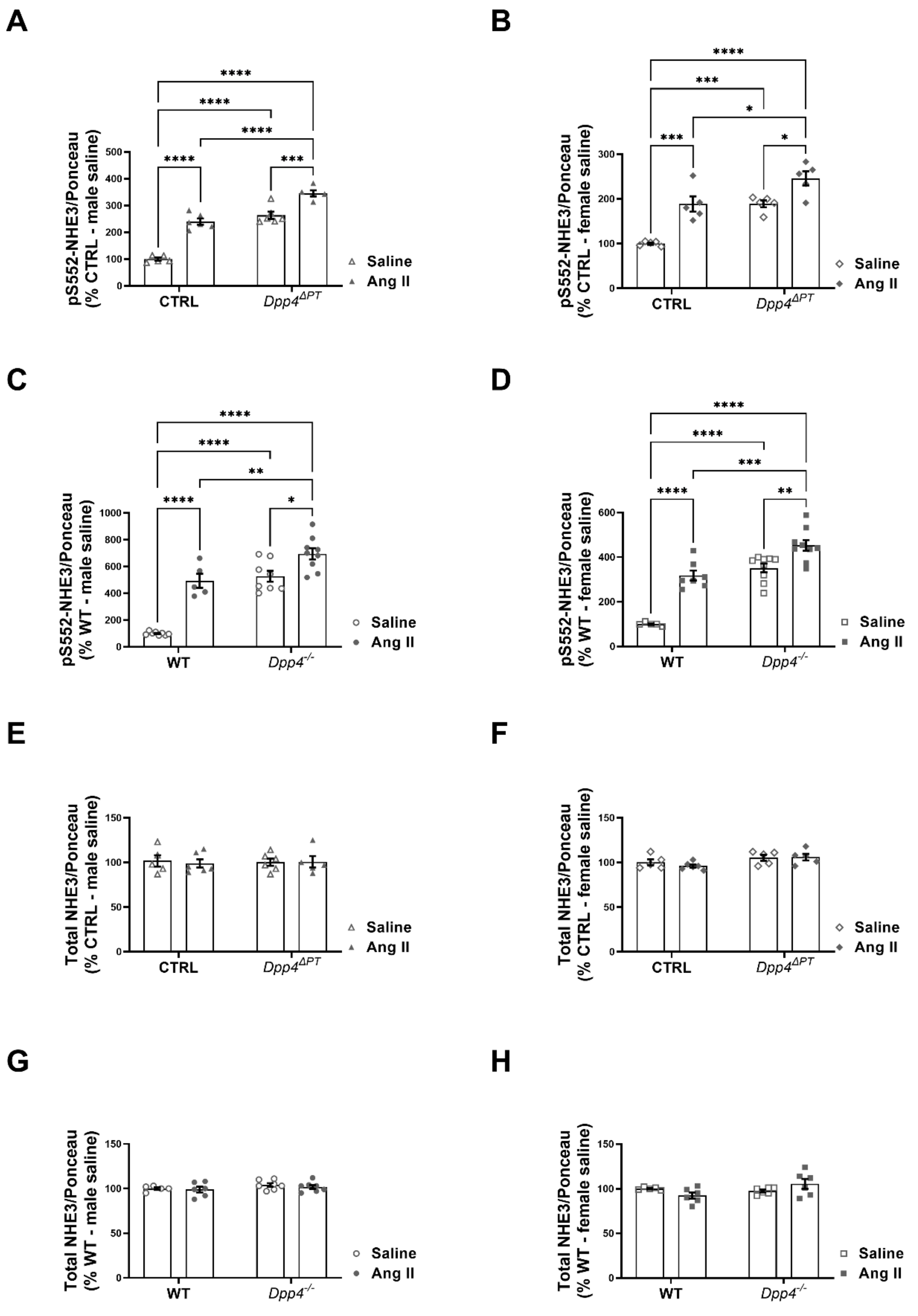


**Supplementary Figure S8 – Influence of acute Ang II-induced blood pressure rise on NHE3 phosphorylation in the kidneys of *Dpp4*$^{\Delta PT}$ and *Dpp4*$^{-/-}$ mice.** Graphical representation of the densitometric analysis of the immunoblotting performed with antibodies against pS552-NHE3 and NHE3 in kidney homogenates from *Dpp4*$^{\Delta PT}$ and *Dpp4*$^{-/-}$ mice. The The densitometry results were normalized to Ponceau (~42 kDa). Data expressed as mean ± SEM, with dots representing the % of CTRL or WT male or female for each animal. Statistical analysis was performed using two-way ANOVA followed by Tukey's post-test. *P < 0.05, **P < 0.01, ***P < 0.001, ****P < 0.0001.

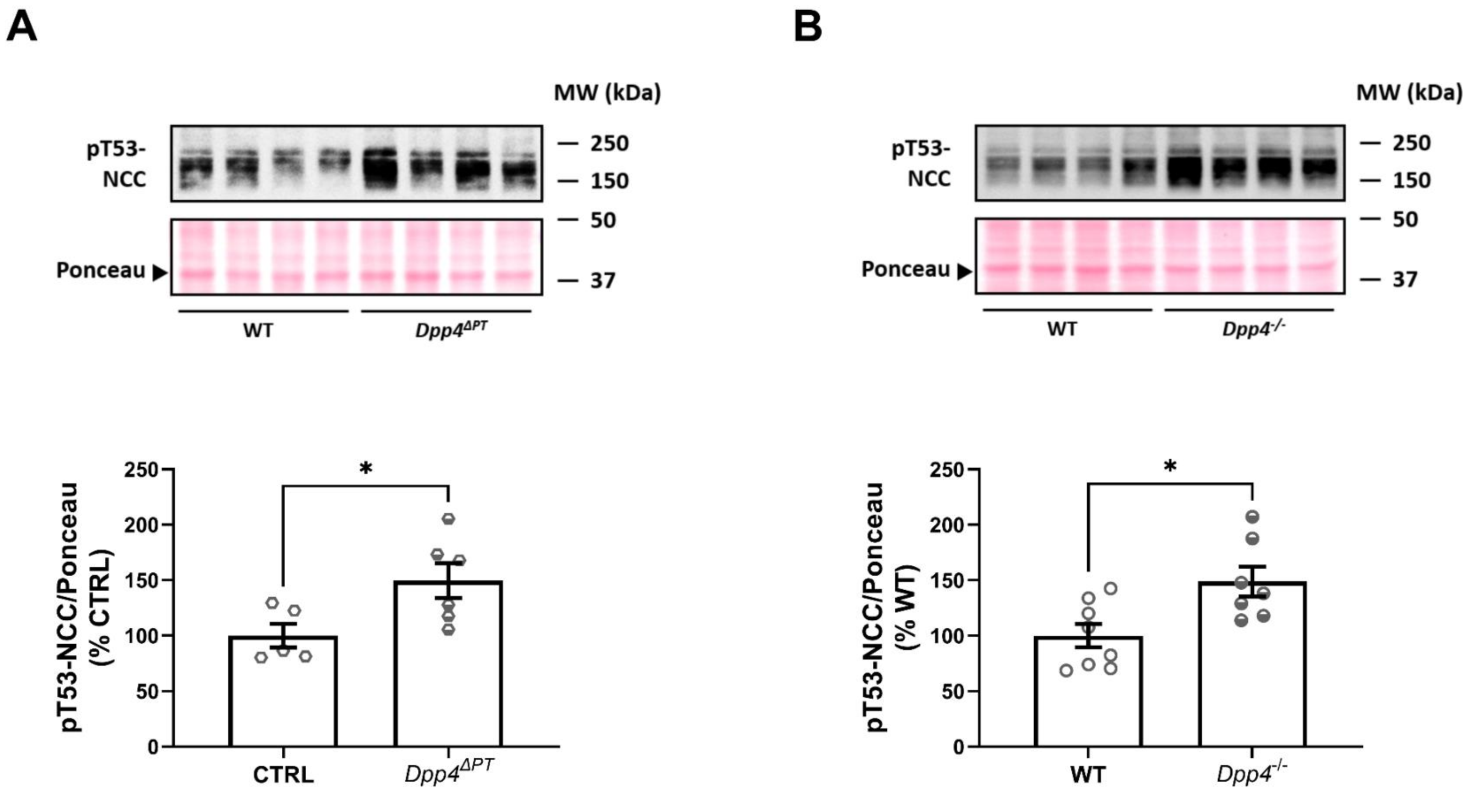


**Supplementary Figure S9 – NCC phosphorylation at threonine 53 (pT53-NCC) in male *Dpp4*$^{\Delta PT}$ and *Dpp4*$^{-/-}$ mice.** pT53-NCC abundance was analyzed by immunoblotting in kidney homogenates from male **(A)** CTRL and *Dpp4*$^{\Delta PT}$ and **(B)** WT and *Dpp4*$^{-/-}$ mice. Densitometry results were normalized to Ponceau staining (~42 kDa). Data expressed as mean ± SEM, with dots representing the % of CTRL or WT male for each animal. Statistical analysis was performed using Student's t-test. *$P < 0.05$.

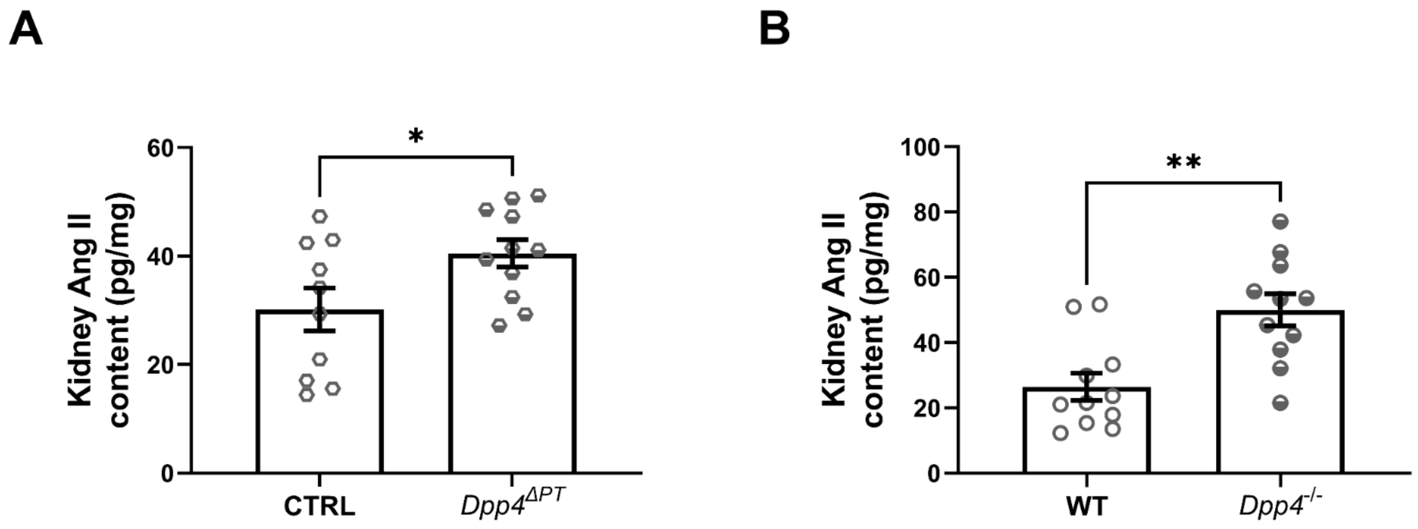


**Supplementary Figure S10 - Renal angiotensin II (Ang II) content in *Dpp4*$^{ΔPT}$ and *Dpp4*$^{-/-}$ mice.** Renal (Ang II) levels were measured by enzyme-linked immunosorbent assay in **(A)** male CTRL and *Dpp4*$^{ΔPT}$ mice and **(B)** male WT and *Dpp4*$^{-/-}$ mice. Data expressed as mean ± SEM, with dots representing individual measurements. Statistical analysis was performed using Student's t-test. *P < 0.05, **P < 0.01.

### FULL UNEDITED GELS

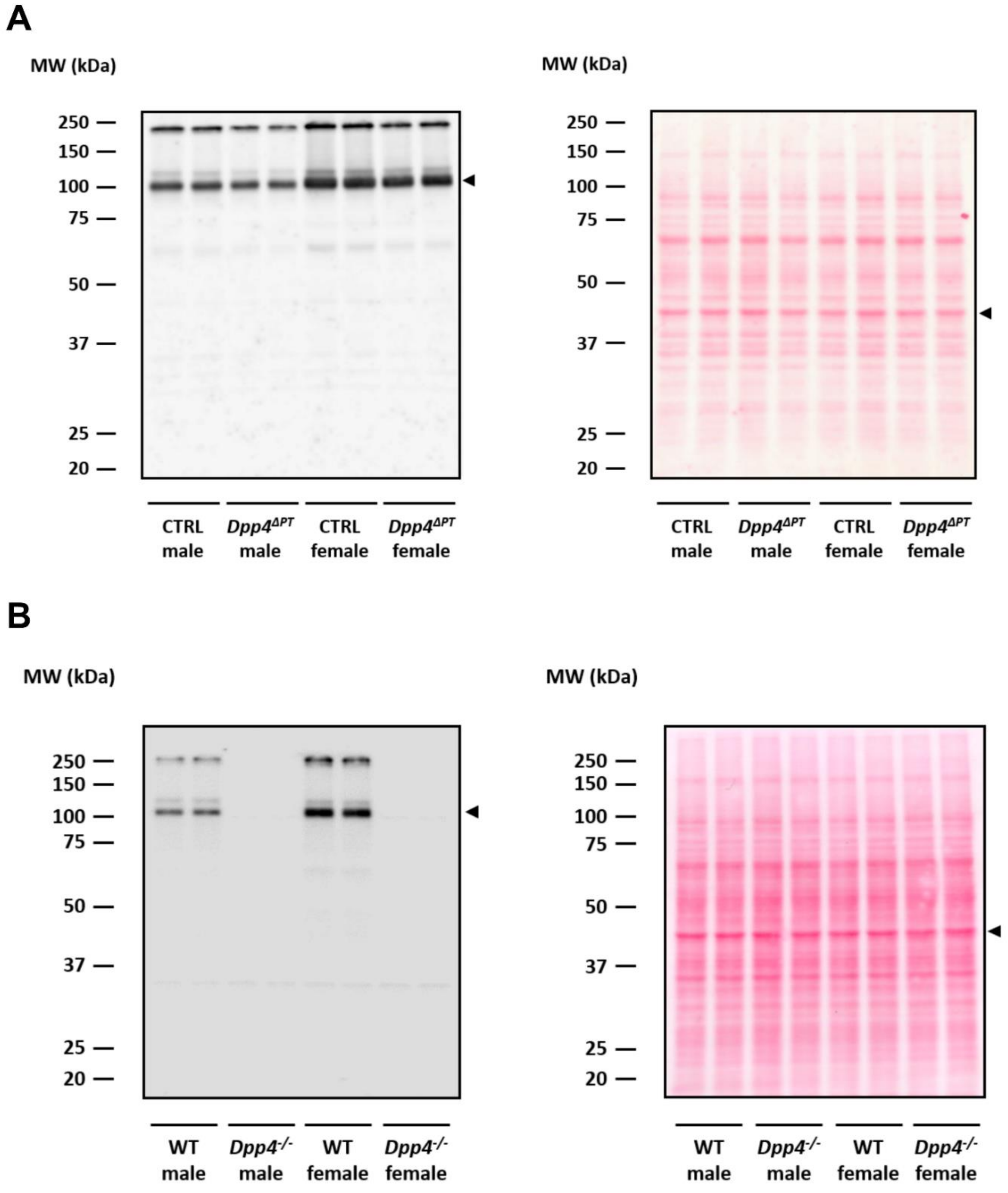


**Supplementary Figure S11.** Full unedited gel for Figure 1. **(A)** Left: Anti-DPP4 blot; Right: Ponceau staining (Figure 1A). **(B)** Left: Anti-DPP4 blot; Right: Ponceau staining (Figure 1B).

**A**

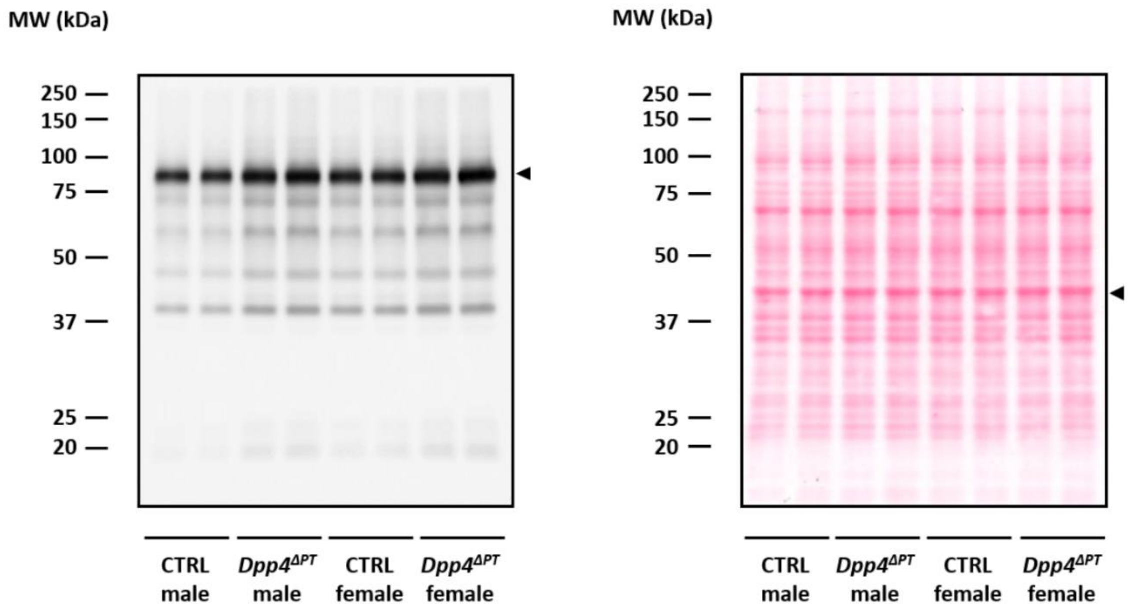
MW (kDa)
250
150
100
75
50
37
25
20
CTRL male
Dpp4ΔPT male
CTRL female
Dpp4ΔPT female
MW (kDa)
250
150
100
75
50
37
25
20
CTRL male
Dpp4ΔPT male
CTRL female
Dpp4ΔPT female

**B**

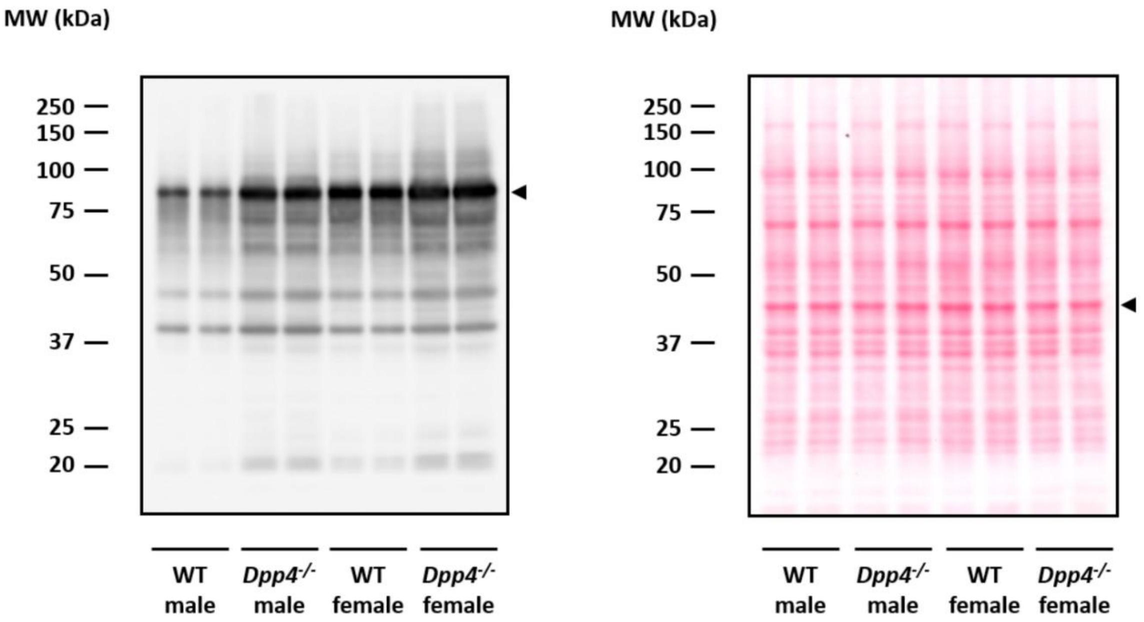
MW (kDa)
250
150
100
75
50
37
25
20
WT male
Dpp4-/- male
WT female
Dpp4-/- female
MW (kDa)
250
150
100
75
50
37
25
20
WT male
Dpp4-/- male
WT female
Dpp4-/- female

C

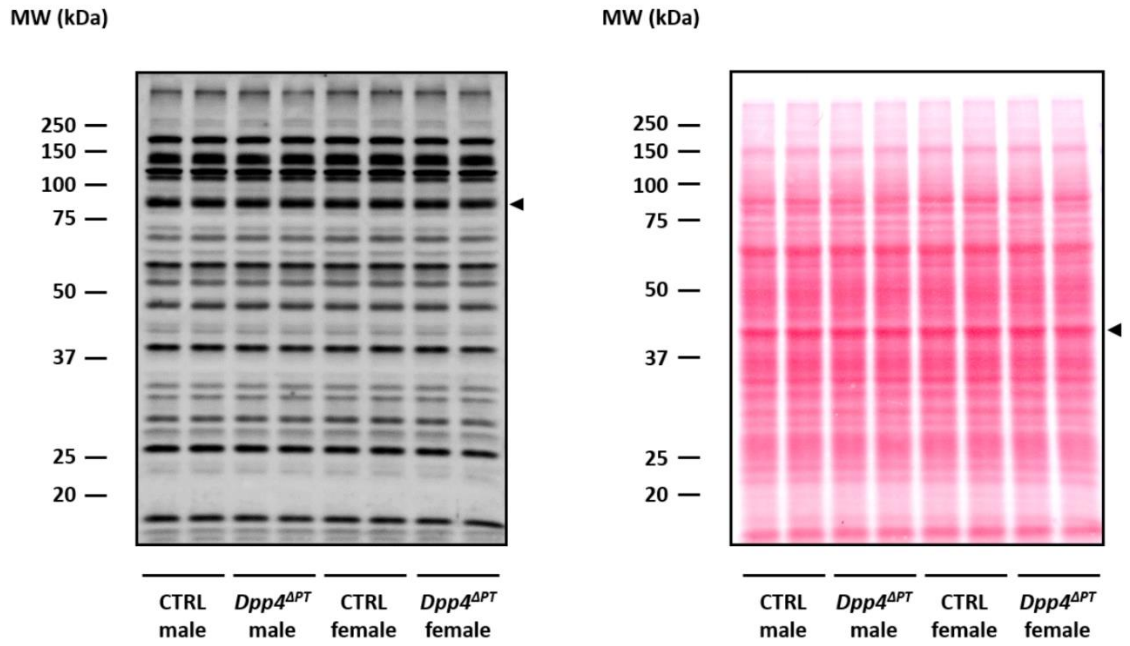


D

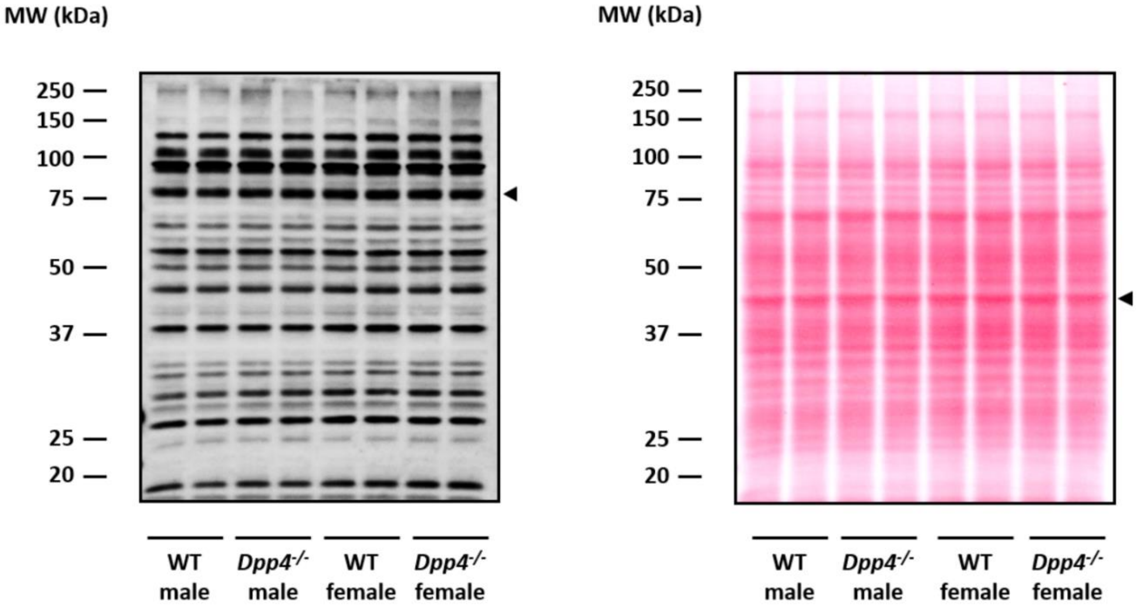


**Supplementary Figure S12.** Full unedited gel for Figure 3. **(A)** Left: Anti-pS552-NHE3 blot; Right: Ponceau staining (Figure 3A). **(B)** Left: Anti-pS552-NHE3 blot; Right: Ponceau staining (Figure 3B). **(C)** Left: NHE3 blot; Right: Ponceau staining (Figure 3A). **(D)** Left: NHE3 blot; Right: Ponceau staining (Figure 3B).

A

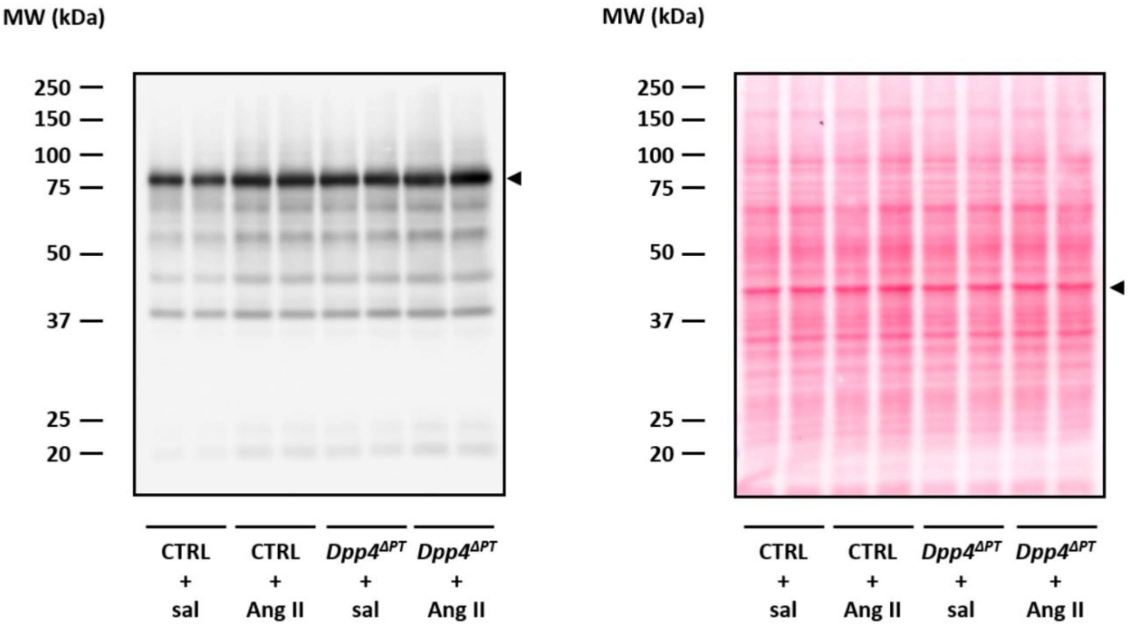
MW (kDa)
250
150
100
75
50
37
25
20
CTRL + sal
CTRL + Ang II
Dpp4^ΔPT + sal
Dpp4^ΔPT + Ang II
MW (kDa)
250
150
100
75
50
37
25
20
CTRL + sal
CTRL + Ang II
Dpp4^ΔPT + sal
Dpp4^ΔPT + Ang II

B

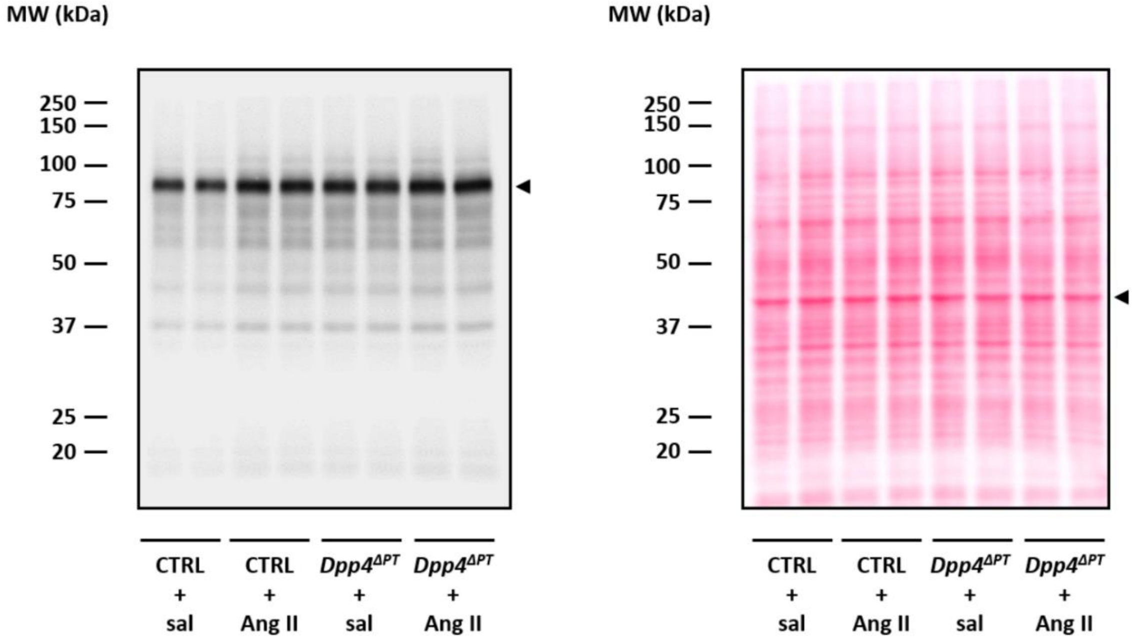
MW (kDa)
250
150
100
75
50
37
25
20
CTRL + sal
CTRL + Ang II
Dpp4^ΔPT + sal
Dpp4^ΔPT + Ang II
MW (kDa)
250
150
100
75
50
37
25
20
CTRL + sal
CTRL + Ang II
Dpp4^ΔPT + sal
Dpp4^ΔPT + Ang II

C

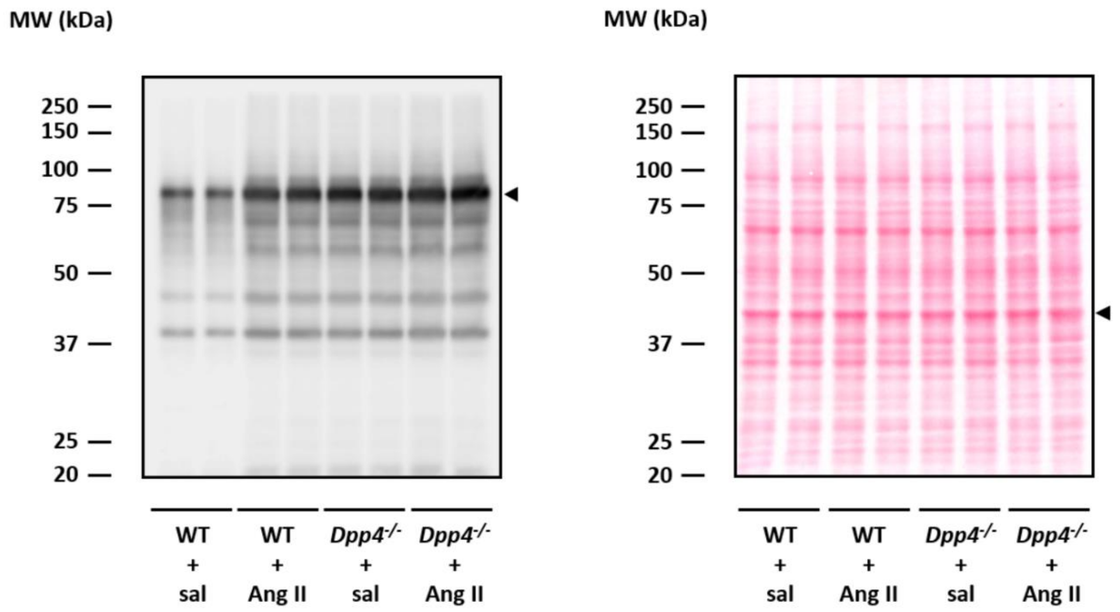
MW (kDa)
250
150
100
75
50
37
25
20
MW (kDa)
250
150
100
75
50
37
25
20
WT + sal
WT + Ang II
Dpp4-/- + sal
Dpp4-/- + Ang II
WT + sal
WT + Ang II
Dpp4-/- + sal
Dpp4-/- + Ang II

D

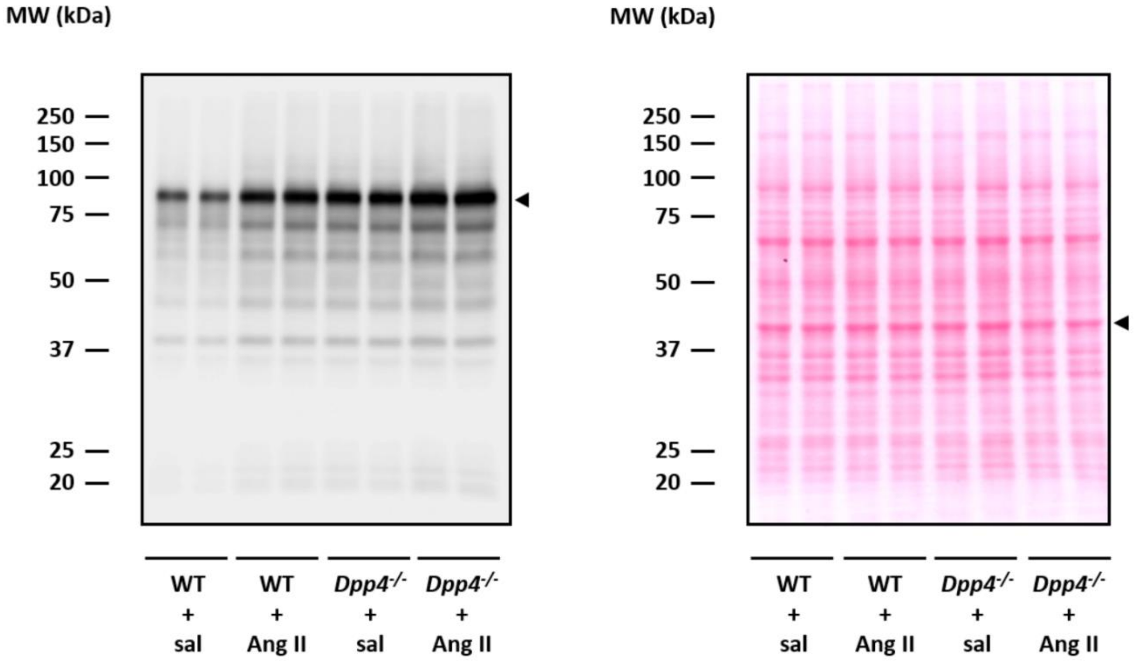
MW (kDa)
250
150
100
75
50
37
25
20
MW (kDa)
250
150
100
75
50
37
25
20
WT + sal
WT + Ang II
Dpp4-/- + sal
Dpp4-/- + Ang II
WT + sal
WT + Ang II
Dpp4-/- + sal
Dpp4-/- + Ang II

E

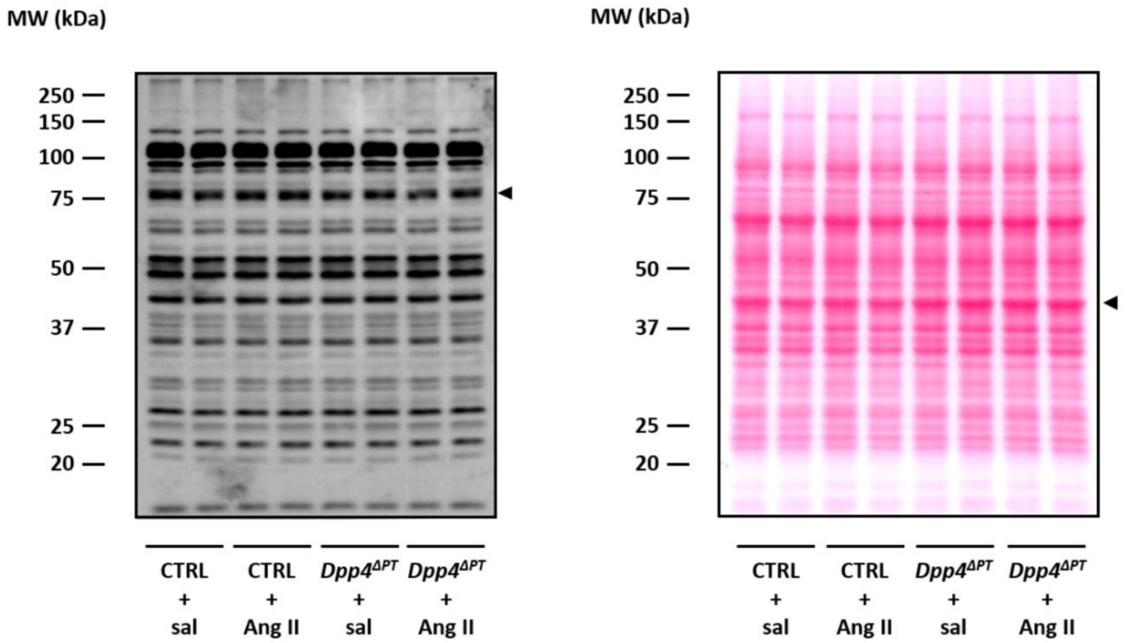
MW (kDa)
250
150
100
75
50
37
25
20
CTRL + sal
CTRL + Ang II
Dpp4ΔPT + sal
Dpp4ΔPT + Ang II
MW (kDa)
250
150
100
75
50
37
25
20
CTRL + sal
CTRL + Ang II
Dpp4ΔPT + sal
Dpp4ΔPT + Ang II

F

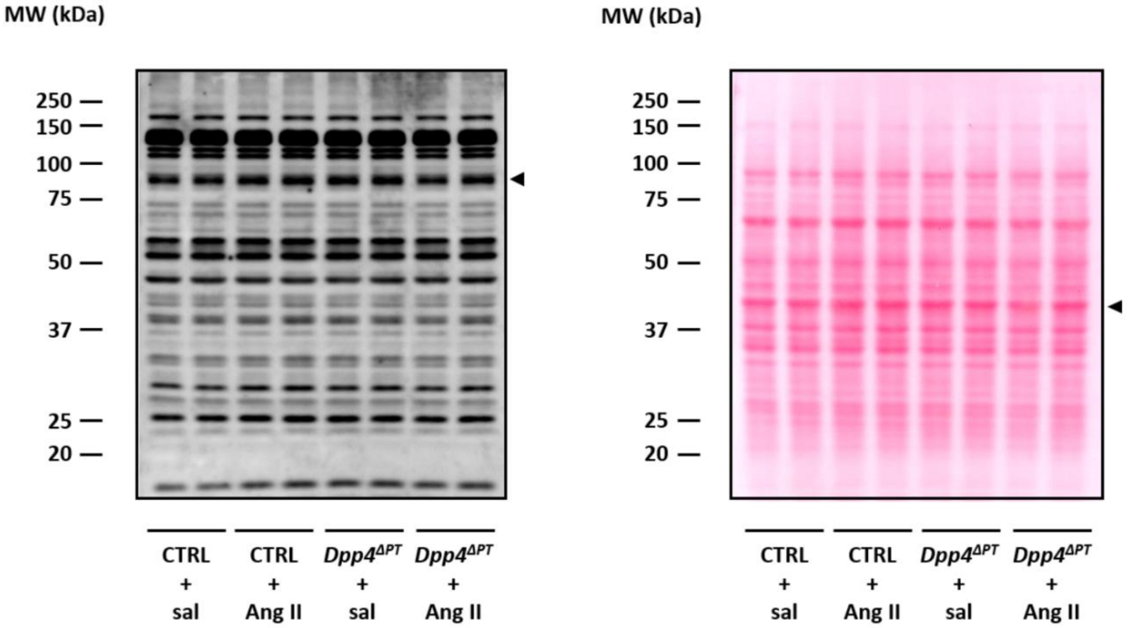
MW (kDa)
250
150
100
75
50
37
25
20
CTRL + sal
CTRL + Ang II
Dpp4ΔPT + sal
Dpp4ΔPT + Ang II
MW (kDa)
250
150
100
75
50
37
25
20
CTRL + sal
CTRL + Ang II
Dpp4ΔPT + sal
Dpp4ΔPT + Ang II

G

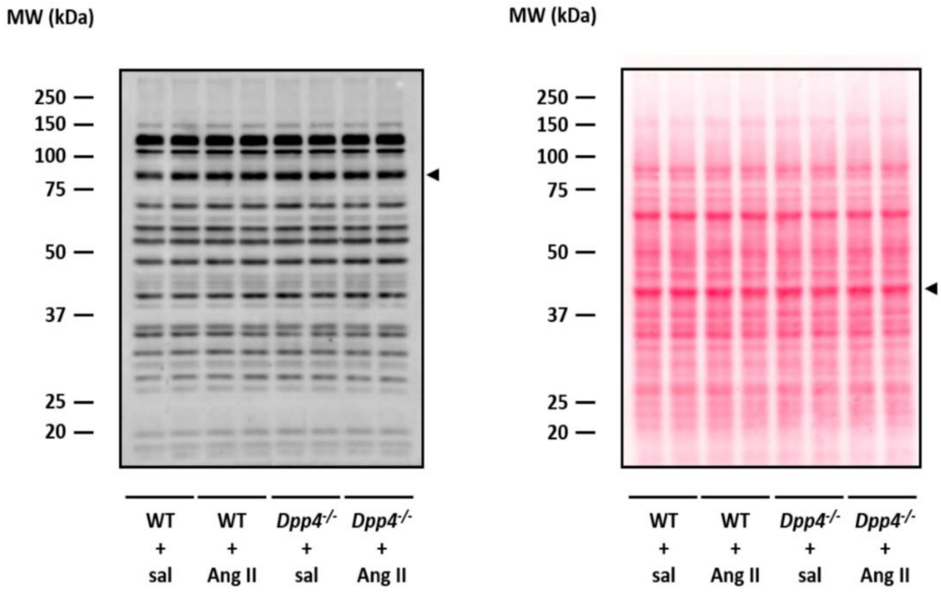


H

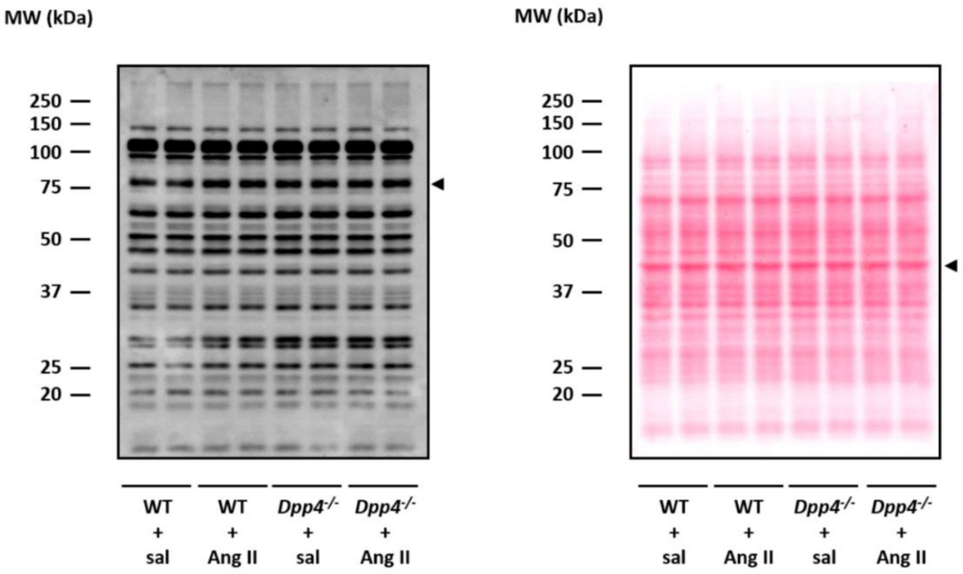


**Supplementary Figure S13.** Full unedited gel for Figure 6. **(A)** Left: Anti-pS552-NHE3 blot; Right: Ponceau staining (Figure 6A). **(B)** Left: Anti-pS552-NHE3 blot; Right: Ponceau staining (Figure 6B). **(C)** Left: Anti-pS552-NHE3 blot; Right: Ponceau staining (Figure 6C). **(D)** Left: Anti-pS552-NHE3 blot; Right: Ponceau staining (Figure 6D). **(E)** Left: NHE3 blot; Right: Ponceau staining (Figure 6A). **(F)** Left: NHE3 blot; Right: Ponceau staining (Figure 6B). **(G)** Left: NHE3 blot; Right: Ponceau. **(H)** Left: NHE3 blot; Right: Ponceau staining (Figure 6D).

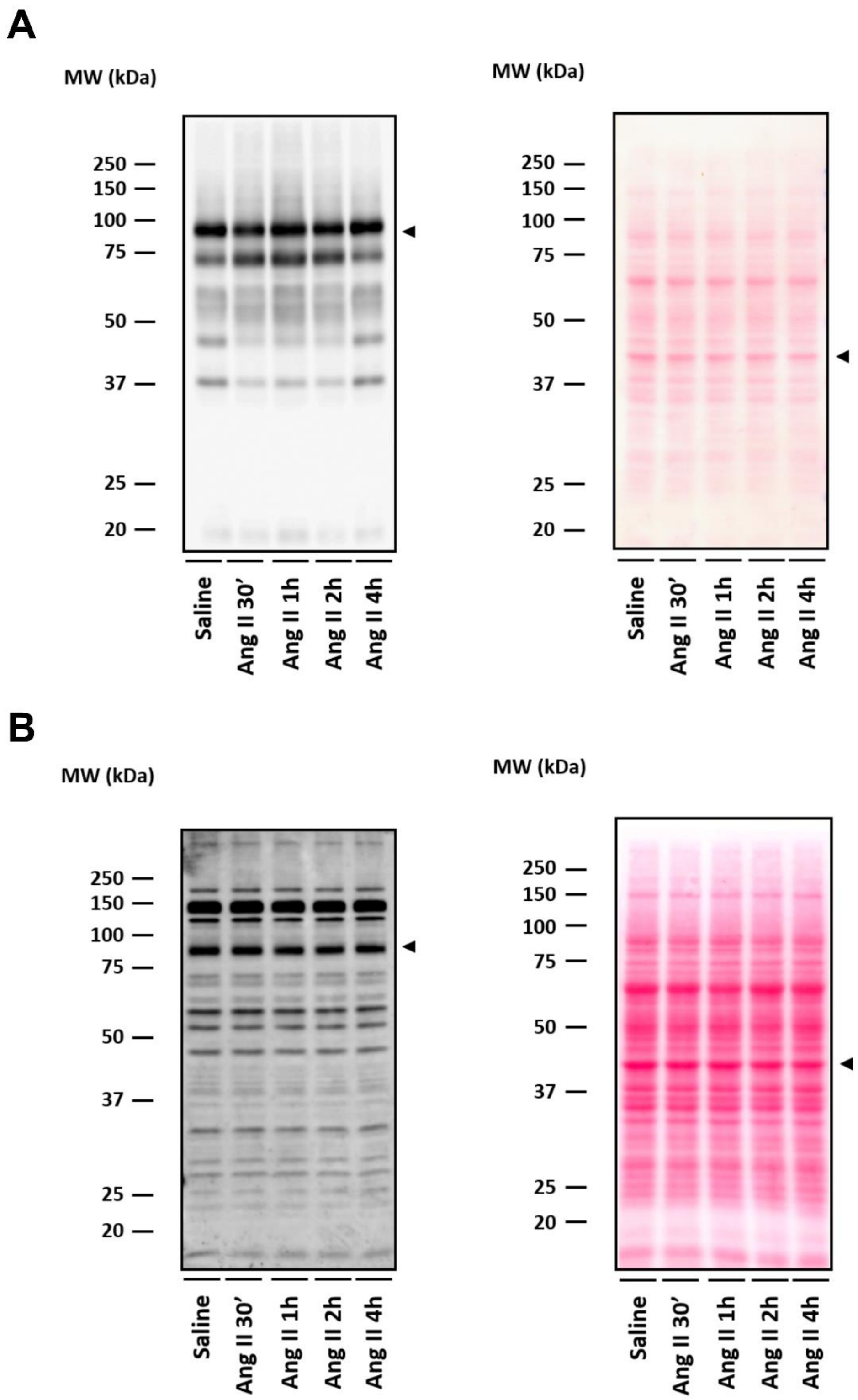


**Supplementary Figure S14.** Full unedited gel for Supplementary Figure S4. **(A)** Left: Anti-pS552-NHE3 blot; Right: Ponceau staining for male WT kidney homogenates (Supplementary Figure S4B). **(B)** Left: NHE3 blot; Right: Ponceau staining (Supplementary Figure S4B).

A

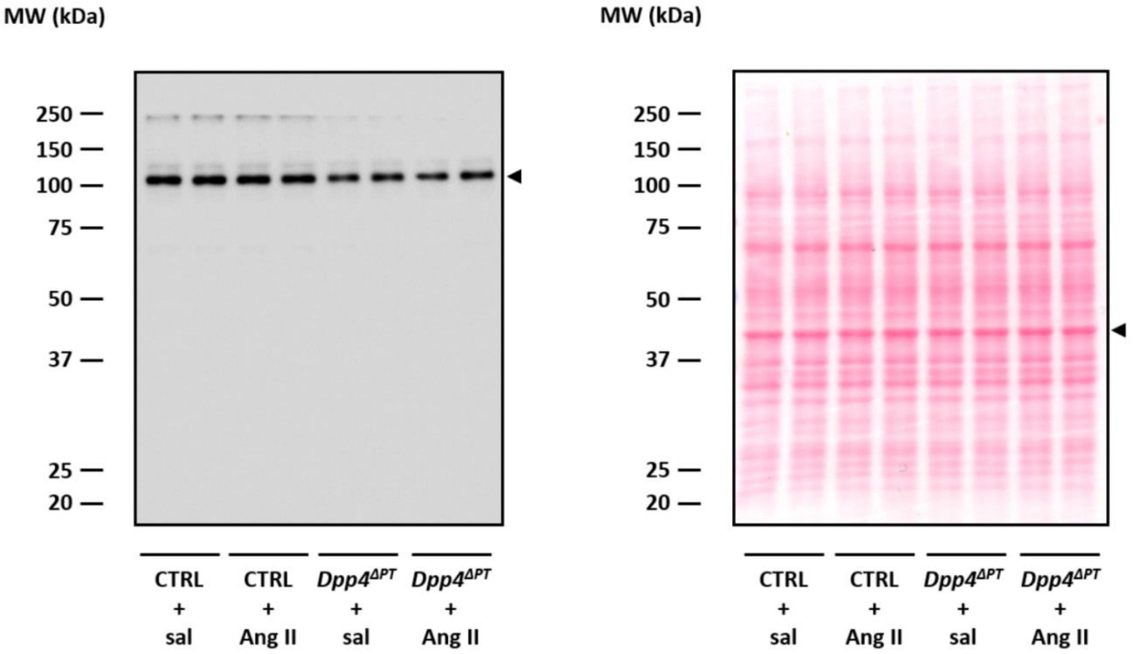
MW (kDa)
250
150
100
75
50
37
25
20
CTRL + sal
CTRL + Ang II
$Dpp4^{\Delta PT}$ + sal
$Dpp4^{\Delta PT}$ + Ang II
MW (kDa)
250
150
100
75
50
37
25
20
CTRL + sal
CTRL + Ang II
$Dpp4^{\Delta PT}$ + sal
$Dpp4^{\Delta PT}$ + Ang II

B

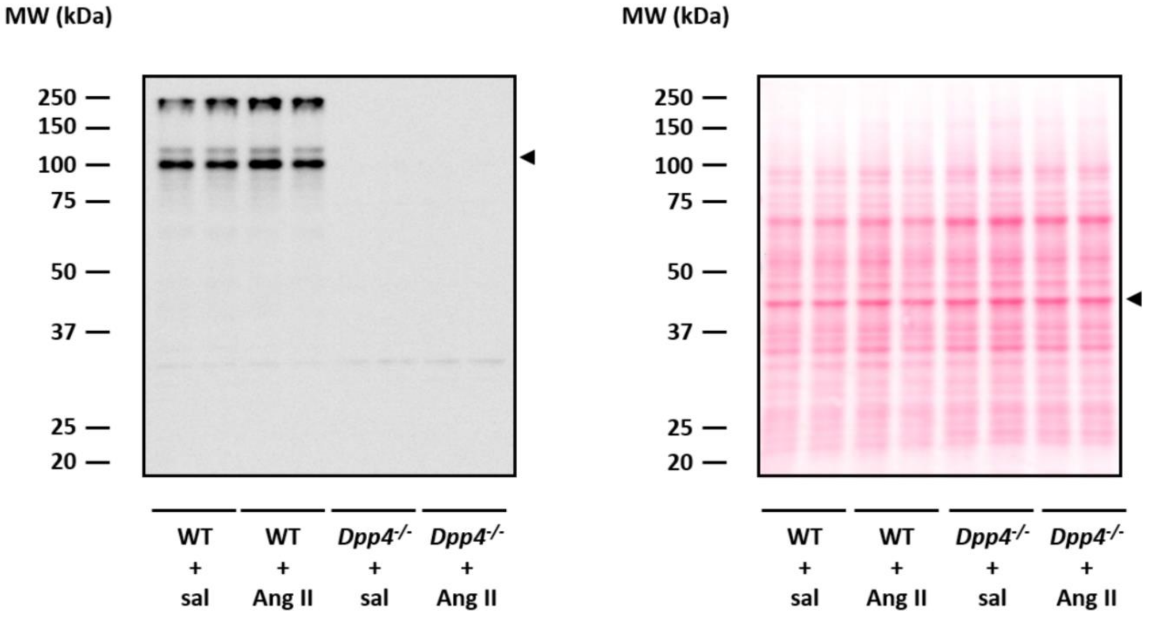
MW (kDa)
250
150
100
75
50
37
25
20
WT + sal
WT + Ang II
$Dpp4^{-/-}$ + sal
$Dpp4^{-/-}$ + Ang II
MW (kDa)
250
150
100
75
50
37
25
20
WT + sal
WT + Ang II
$Dpp4^{-/-}$ + sal
$Dpp4^{-/-}$ + Ang II

C

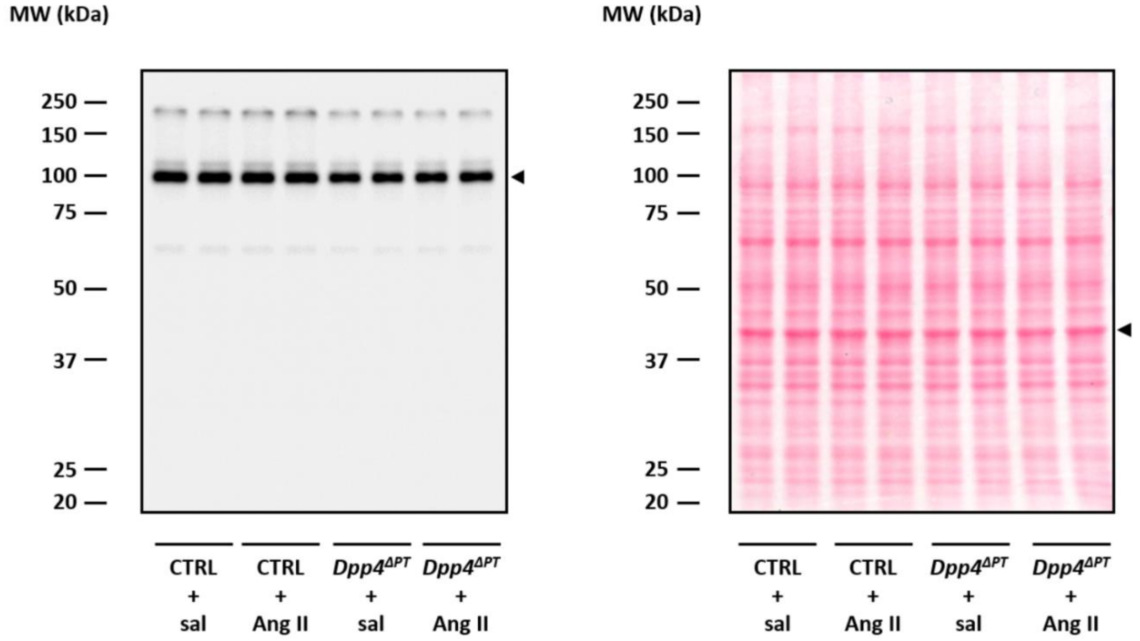


D

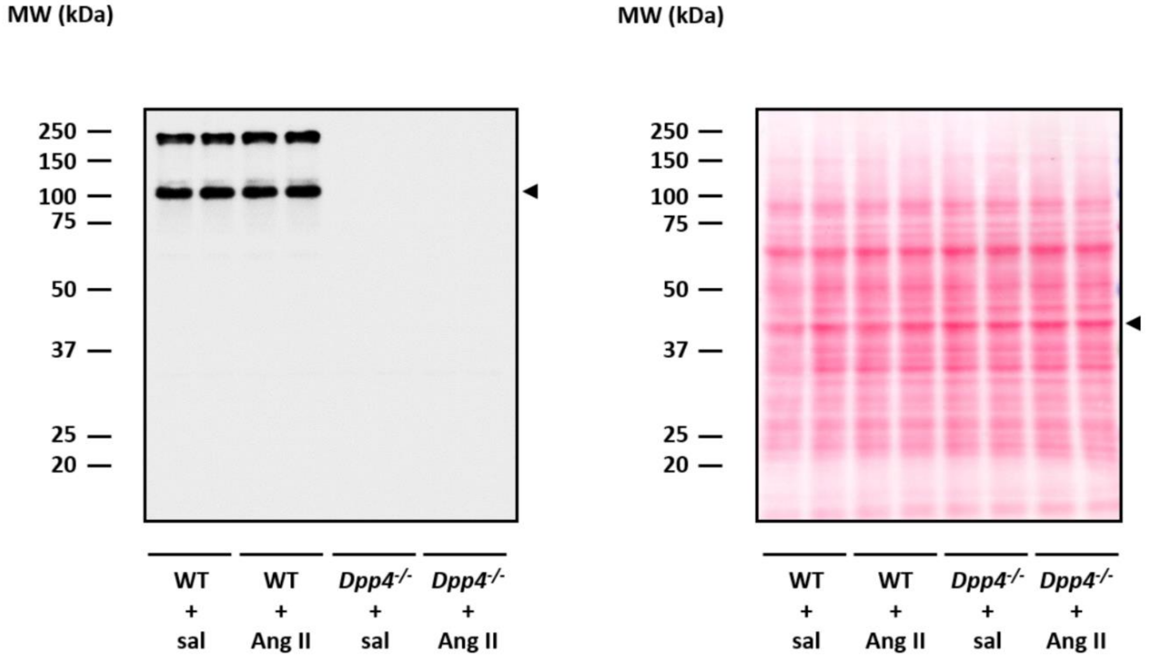


**Supplementary Figure S15.** Full unedited gel for Supplementary Figure S5. **(A)** Left: DPP4 blot; Right: Ponceau staining (Supplementary Figure S5A). **(B)** Left: DPP4 blot; Right: Ponceau staining (Supplementary Figure S5B). **(C)** Left: DPP4 blot; Right: Ponceau staining (Supplementary Figure S5C). (D) Left: DPP4 blot; Right: Ponceau staining (Supplementary Figure S5D).

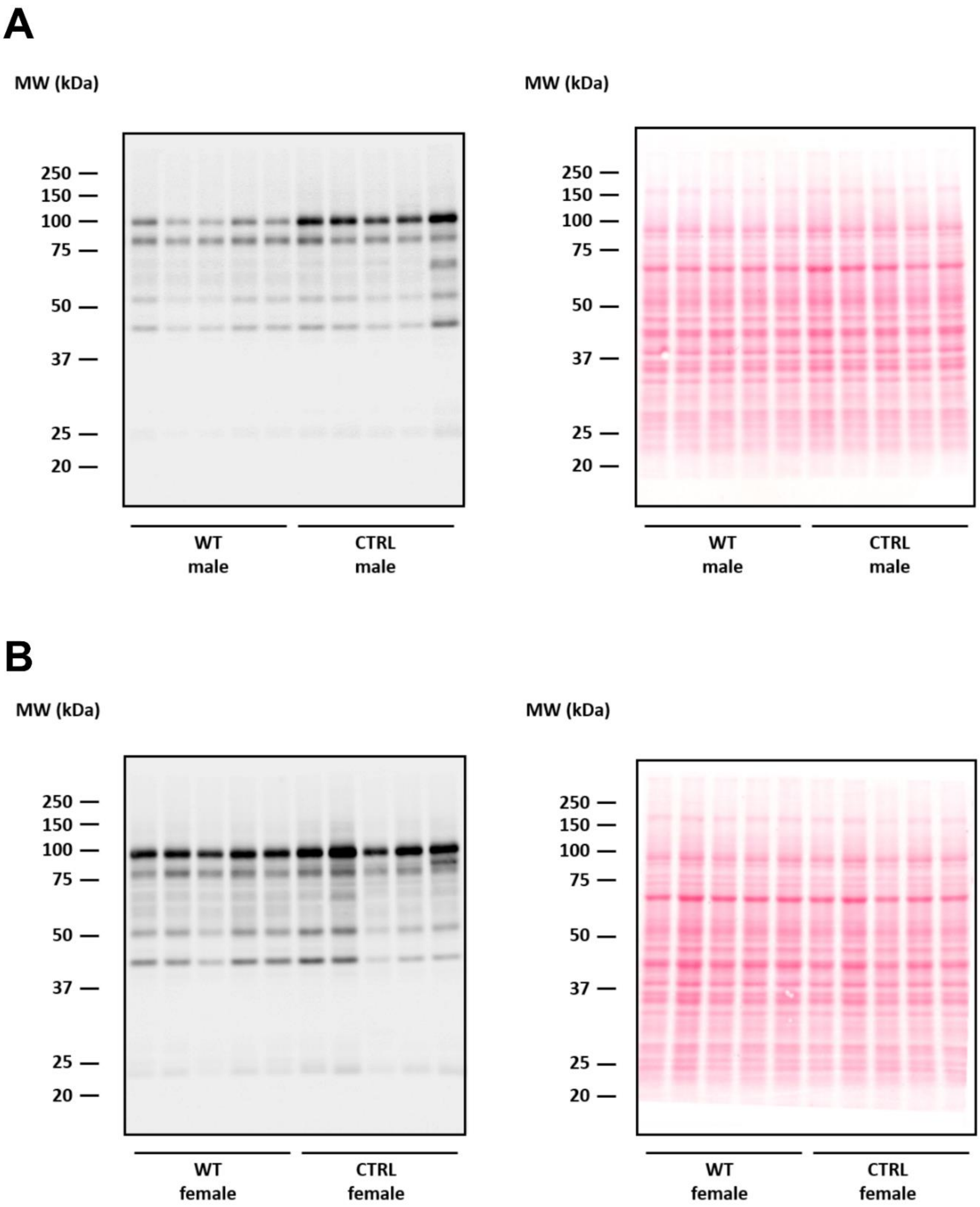


**Supplementary Figure S16 -** Full unedited gel for Supplementary figure S8. **A)** Left panel: pS552-NHE3 and Right panel: ponceau staining, in CTRL and WT male mice kidneys homogenates (Supplementary figure S8A). **B)** Left panel: pS552-NHE3 and Right panel: ponceau staining, in CTRL and WT female mice kidneys homogenates (Supplementary figure S8B).

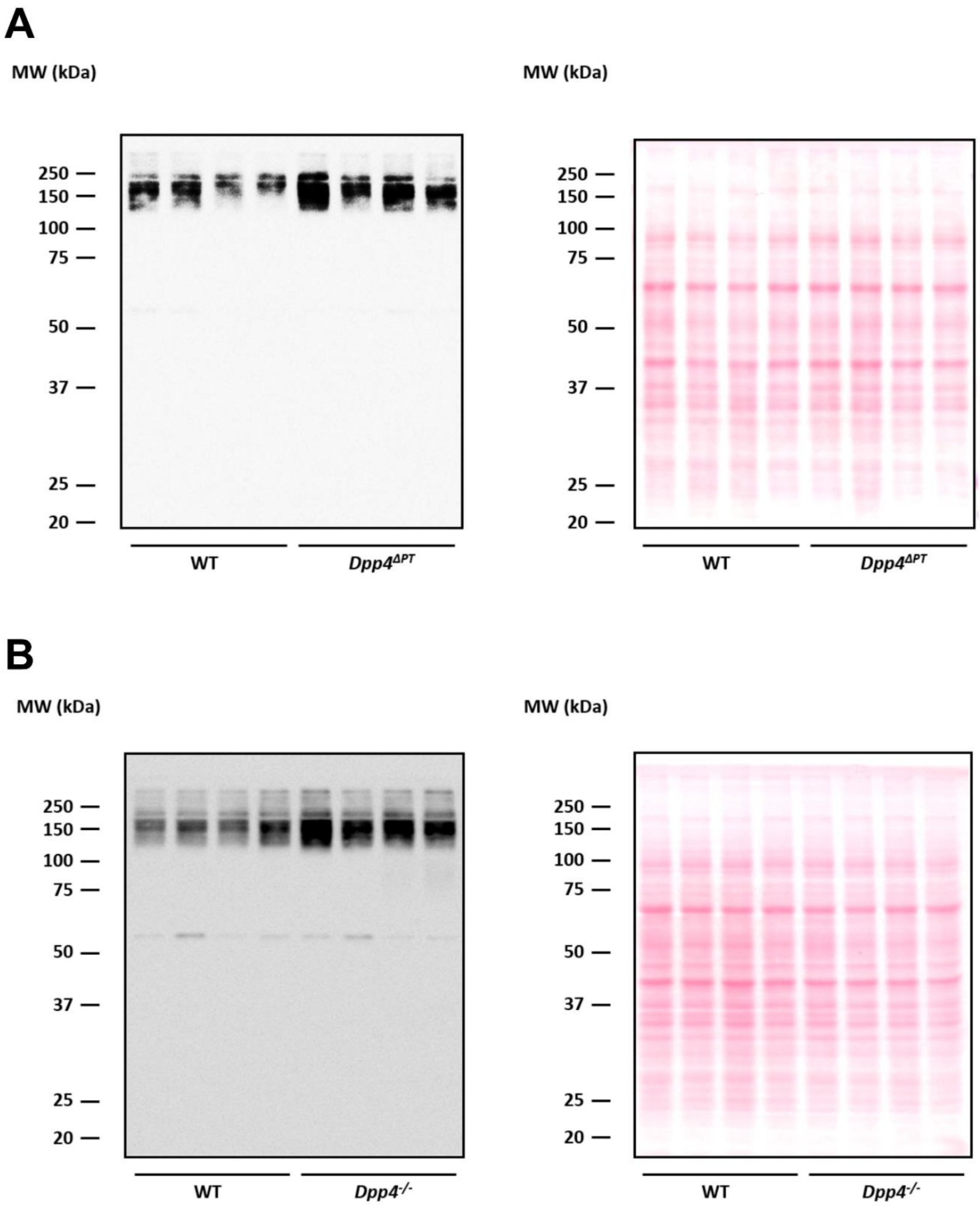


**Supplementary Figure S17.** Full unedited gel for Supplementary Figure S9. **(A)** Left: pT53-NCC blot; Right: Ponceau staining (Supplementary Figure S9A). **(B)** Left: pT53-NCC blot; Right: Ponceau staining (Supplementary Figure S9B).
